# Lipid nanoparticle co-delivery of mRNA and a small molecule drug for oral cancer chemoimmunotherapy

**DOI:** 10.1101/2025.09.26.678832

**Authors:** Marshall S. Padilla, Jacqueline J. Li, Qunzhou Zhang, Korey Patwari, Shihong Shi, Hannah M. Yamagata, Ryann A. Joseph, Briyanna N. Hymms, Sridatta V. Teerdhala, Mohamad-Gabriel Alameh, Drew Weissman, Anh D. Le, Michael J. Mitchell

## Abstract

Oral squamous cell carcinoma (OSCC) represents 90% of all head and neck cancers. Despite decades of research, the 5-year survival rate is 50%, and strikingly, the overall incidence rate is projected to increase by 30% in the next ten years, which will result in a sharp increase in mortality. Two fundamental aspects of OSCC are that it progresses via the inactivation and mutation of tumor suppressor (TS) genes and has a “cold” tumor immune microenvironment (TIME). A major barrier in the treatment of OSCC is the lack of novel therapies clinicians have at their disposal that are designed to disrupt tumor progression by reshaping the cold tumors into inflammatory “hot” tumors. To overcome these obstacles, we employed a lipid nanoparticle (LNP) that co-encapsulates p53 mRNA and the small molecule ciclopirox (CPX). We demonstrate that both drugs have innate chemotherapeutic properties by facilitating caspase activation. Moreover, these therapies can create a less immunosuppressive TIME in part by repolarizing tumor-associated macrophages (TAMs) to M1-like phenotypes. When formulated together, our platform provides an all-in-one approach for OSCC, effective in both p53-therapy-susceptible and p53-therapy-resistant models. Additionally, this work provides a template for a delivery platform capable of tackling multiple mechanisms of OSCC progression and survival.

## Introduction

Head and neck squamous cell carcinoma (HNSCC) is the sixth most common form of cancer in the world, where ~450,000 people die from this disease each year, with oral squamous cell carcinoma (OSCC) representing over 90% of HNSCC^1^. Unfortunately, the incidence rate of OSCC is projected to increase by 30% by the end of the decade. Standard of care for oral cancer involves ablative surgery and radiation, and although these therapies are effective, 50% of patients experience difficulty speaking and swallowing as a result of these treatments. or recurrent or metastatic tumors, decades-old chemotherapies such as cisplatin underscore the lack of advancement in this field. Although immunotherapies are promising, they have yet to significantly improve overall survival due to low response rates^2^. To advance patient care, it is pertinent to develop therapies that overcome the specific barriers in OSCC. This includes the large population of tumor associated macrophages (TAMs) in the oral tumor immune microenvironment (TIME) that strongly contribute to cancer immune evasion. Additionally, oral tumors have a high mutational rate of intrinsic tumor suppressor genes (TSGs), including TP53, that leads to their inactivation and contribution to tumor development and progression through regulating DNA repair, cell proliferation, apoptosis, and even the cancer immune response^3,4^.

To address the multitude of mechanisms that cause cancer progression, it has become more common to co-administer therapeutics that target different cancer axes^5,6^. While effective, it can lead to severe toxicity for the patient due to the combined side effects of each drug. Co-formulation into a single drug delivery vehicle represents an exciting alternative as both therapeutics can be simultaneously loaded and administered. Drug delivery vehicles also offer cargo protection and can be tuned to target tumors directly, limiting off-target accumulation^7^. Still, if the therapeutics have different chemical characteristics, such as water solubilities, efficient loading can be difficult. Recently, lipid nanoparticles (LNPs) have emerged as the preeminent vehicle for RNAs, being the FDA-approved carriers for the Pfizer/BioNTech and Moderna COVID-19 mRNA vaccines^8^. LNPs have aqueous and lipid microenvironments, allowing for the encapsulation of hydrophilic and lipophilic drugs. For example, we have demonstrated that LNPs can be co-loaded with Bcl-2 siRNA and doxorubicin for synergistic cancer therapy^9^. Even though there has been research in optimizing LNPs for OSCC, there is no research on co-delivery of multiple therapeutics with LNPs for OSCC therapy^10,11^.

Here, we construct an LNP platform that co-encapsulates RNA and a small molecule drug to reduce tumor viability and polarize the TME. For the RNA component, we deliver p53 mRNA as p53 is underexpressed or mutated in over 70% of OSCC cases^12^. Several p53 restoration therapies have been developed for hepatocarcinoma and non-small cell lung cancer; however, no p53 mRNA therapies have been utilized for OSCC^13,14^. p53 expression mediates apoptosis in cancer cells, and recent studies have found that increased p53 expression can halt M2-like polarization in macrophages^15^. However, p53 therapies have failed in clinical trials due to mediocre performance and the toxicity of viral vector delivery vehicles that are the current standard for tumor suppressor replacement therapy. Although OSCC tumors can be broadly classified as p53-deficient, the cell populations have heterogenous expression of wild type p53 and mutated p53 species. Thus, the purpose of this study is to co-deliver p53 mRNA and ciclopirox (CPX), an FDA-approved anti-fungus medication that has been repurposed as a chemotherapeutic in bladder cancer, pancreatic cancer, and colorectal cancer in preclinical studies^16,17^.

Both drugs require a drug delivery carrier as CPX has poor water solubility and p53 mRNA cannot cross cell membranes due its large size, negative charge, and susceptibility for degradation by nucleases^18^. To achieve potent transfection of both drugs, we co-encapsulated them into a single LNP formulation (**Figure 1A**). CPX is soluble in ethanol and thus can be incorporated into the ethanolic phase of the LNP formulation process that also contains the other lipid excipients. The resulting LNP facilitated synergistic cell killing in human and murine OSCC lines, which we determined to be a result of increased caspase activity. Moreover, we determined that both p53 and CPX can repolarize TAMs to a more M1-like phenotype in OSCC, although the relative effect of each therapeutic depends on the cell type (**Figure 1B–D**). We then demonstrate that the combination therapy allows for the reduction of tumor burden in models where one therapy is ineffective. This highlights that this platform can be implemented for a wider range of OSCC tumors in place of traditional monotherapies.

**Figure 1.**
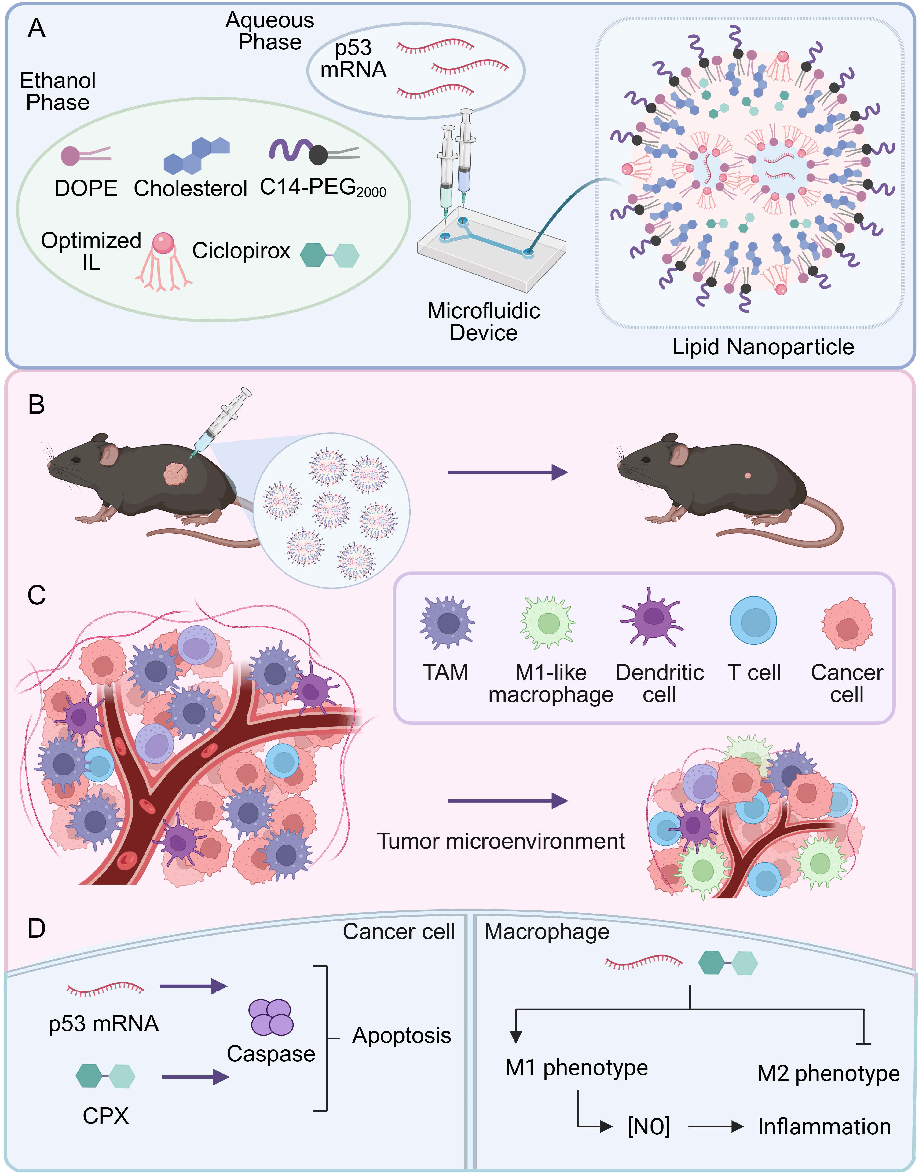
Overview of LNP co-delivery platform. **A**, LNPs are formulated via a microfluidic device, combining an aqueous mRNA phase with an ethanolic phase containing the lipid excipients. **B–C**, LNP treatment delays tumor growth by **(B)** killing tumor cells and **(C)** altering the TIME via polarization of TAMs to M1-like macrophages. **D**, Cancer cell killing is mediated by caspase pathways while macrophage polarization involves the expression of M1-like markers and increased nitric oxide production.

## Results

### Optimization of a potent lipid nanoparticle for oral squamous cell carcinoma

LNPs are formulated using an ethanol phase that dissolves lipid components and a citrate phase that contains the mRNA. The lipids include a phospholipid, cholesterol, PEGylated lipid, and an ionizable lipid. The ionizable lipid is central to the efficacy of the LNP due to its structure that amalgamates amines and hydrophobic tails. Amines allow for switchable cationic charge, whereas the tails endow the lipid with hydrophobicity. As such, ionizable lipids protonate in the citrate buffer, allowing the molecules to electrostatically bind to negatively charged mRNA, enwrapping it in the hydrophobic tails, and enabling it to be encapsulated in the LNP. After formulation, the LNPs are dialyzed against phosphate buffer saline (PBS) to neutralize the charge, as particles with permanent positive charges can induce toxicity^19^. Once LNPs enter cells, they proceed through the low pH endo-lysosomal pathway, allowing the ionizable lipids to regain their positive charge. The positively charged ionizable lipids then disrupt the endosomes and lysosomes and facilitate mRNA escape into the cytosol to begin translation^20^. Since ionizable lipids play a pivotal role in LNP delivery, it is important to tailor their structure as different lipids are optimal for certain biological environments^21,22^.

We have recently developed a class of ionizable lipids with terminal branching that enhance endosomal escape leading to more efficacious mRNA delivery^23^. These Branched ENdosomal Disruptor (BEND) lipids were synthesized by reacting a linear or branched alkyl epoxide with a polyamine core via an S_N_2 reaction (**Figure 2A, Supplementary Figure 1**). Each ionizable lipid varied in its total alkyl length as well as branching type, with the latter including non-branched or linear, isopropyl, tert-butyl, and sec-butyl structures, with sixteen ionizable lipids being synthesized in total. The ionizable lipids were then formulated into LNPs with firefly luciferase (FLuc) mRNA using a molar ratio of 35:16:46.5:2.5 for the ionizable lipid, 1,2-dioleoyl-sn-glycero-3-phosphoethanolamine (DOPE), cholesterol, and 14:0 PEG2000 PE, respectively. LNPs were prepared at a 10:1 ionizable lipid to mRNA weight ratio using herringbone microfluidic devices^24^. Each of the ionizable lipids were formulated into separate LNPs, with the gold standard, C12-200, prepared as a positive control for a total of seventeen unique LNP formulations. Traditionally, in vitro assays are used to screen LNPs for in vivo applications; however, transfection trends in 2D monolayers often do not correlate with in vivo trends^25^. Instead, we developed a multi-tiered screening approach to identify a lead LNP candidate that has high transfection in oral cancer cells and low off-target mRNA delivery in the liver. For the first step, the LNPs were incubated in CAL-27 cells, a common human tongue squamous cell carcinoma line, at a dose of 20 ng of encapsulated mRNA per 20,000 cells. After 24 h, a luminescence assay (**Figure 2B**) and toxicity assay (**Figure 2C**) were performed to evaluate the efficacy and safety of the LNPs in vitro, respectively. Two LNPs, E10i-494 and E10s-494, had a 2-fold increase in luciferase expression compared to C12-200, while no LNPs had viabilities under 75%. Utilizing previous data on intravenous liver delivery of BEND LNPs, we plotted the liver delivery of each LNP against its transfection in CAL-27 cells and identified four LNPs, C10-494, C12-494, E10i-494, and E10s-494 with low hepatic tropism and potent OSCC delivery (**Figure 2D**). Since LNPs have naturally high accumulation in the liver, identifying formulations with low liver transfection is important for decreasing off-target toxicity. We then created a subcutaneous xenograft tumor model using CAL-27 cells in Nu/J mice. After two weeks, we intratumorally administered the top four LNPs along with C12-200 at a dose of 0.1 mg/kg (**Figure 2E**). After 6 h, the mice received intraperitoneal injections of D-luciferin and were sacrificed. The heart, lungs, liver, kidneys, spleen, and tumor were dissected and imaged for luminescence by an In Vivo Imaging System (IVIS) (**Figure 2F**). E10i-494 emerged as the lead performer, having 50-fold higher tumor luminescence compared to the gold standard C12-200 (**Figure 2G**).

**Figure 2.**
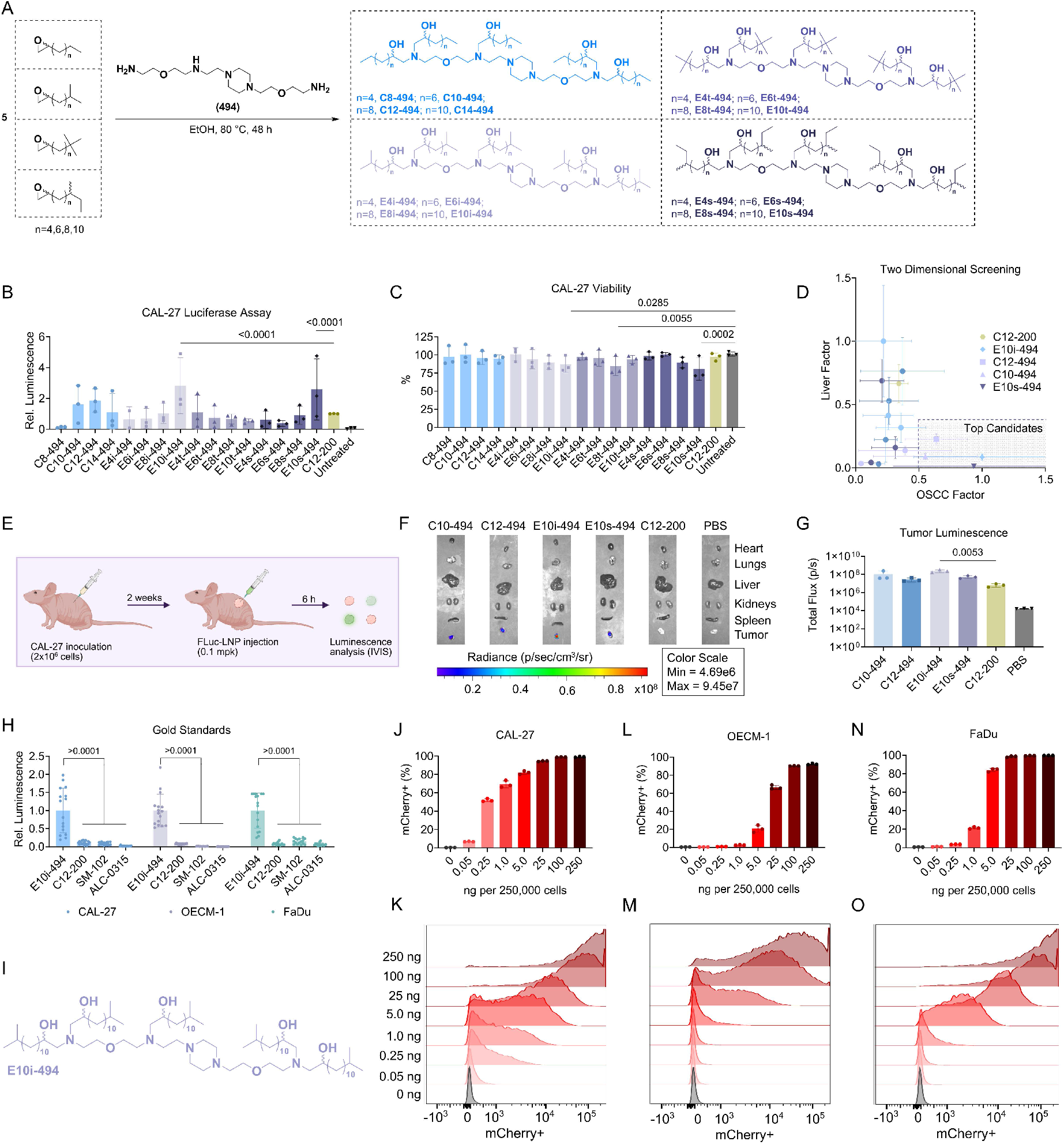
Optimization of LNPs for oral cancer. **A**, Ionizable lipids are synthesized via an S_N_2 reaction between a linear or terminally branched alkyl epoxide and the polyamine, 494. **B–C**, CAL-27 cells were transfected with FLuc-containing LNPs at a dose of 20 ng of mRNA per 20,000 cells. After 24 h, **(B)** luminescence and **(C)** viability were measured. Relative luminescence and viability are reported as mean ± SD of *n* = 3 biological replicates, with each biological replicate consisting of *n* = 4 technical replicates. **D**, CAL-27 luminescence (x-axis) was plotted against liver luminescence after LNP intravenous administration in C57BL/6J mice (y-axis). The liver luminescence data was previously obtained^23^. **E**, Nu/J mice were inoculated with two million CAL-27 cells on the right flank, and after two weeks, LNPs were intratumorally administered at 0.1 mpk. **F–G**, After 6 h, the mice were sacrificed, and organ luminescence images were **(F)** captured and **(G)** quantified utilizing IVIS. Total flux is reported as mean ± SD of *n* = 3. **H**, The top performer, E10i-494, and three gold standards, C12-200, SM-102, and ALC-0315, loaded with FLuc mRNA were dosed in CAL-27, OECM-1, and FaDu cells at 20 ng of mRNA per 20,000 cells. After 24 h, luminescence was quantified. Relative luminescence is reported as mean ± SD of *n* = 16 replicates. **I**, Structure of E10i-494 ionizable lipid. **J–O**, E10i-494 was reformulated with mCherry mRNA transfected in CAL-27, OECM-1, and FaDu cells at increasing dosages per 250,000 cells. After 24 h, the oral cancer cells were analyzed by flow cytometry to measure **(J)** mCherry+ CAL-27 cells, **(K)** CAL-27 fluorescence intensity, **(L)** mCherry+ OECM-1 cells, **(M)** OECM-1 fluorescence intensity, **(N)** mCherry+ FaDu cells, and **(O)** FaDu fluorescence intensity. mCherry positivity is reported as mean ± SD of *n* = 3. Representative data were used for the fluorescence intensity plots. **B–C**, Nested one-way ANOVA with *post hoc* Holm–Šídák correction for multiple comparisons was used to compare **(B)** luminescence against C12-200 and **(C)** viability against untreated cells. **G**, One-way ANOVA with *post hoc* Holm–Šídák correction for multiple comparisons was used to compare tumor luminescence to C12-200. **H**, Two-way ANOVA with *post hoc* Holm–Šídák correction for multiple comparisons was used to compare E10i-494 relative luminescence against individual gold standards for each cell type.

To verify the potency of E10i-494, we compared it against lead gold standards in additional cell lines. E10i-494 was evaluated in CAL-27 cells as well as OECM-1 and FaDu cells, other human OSCC lines, against C12-200, SM-102, and ALC-0315. SM-102 and ALC-0315 are the LNPs utilized in the mRNA COVID-19 vaccine formulations for Moderna and Pfizer/BioNTech, respectively. When dosed at 20 ng per 20,000 cells, E10i-494 outperformed all gold standards in all three cells line by 2–5 fold (**Figure 2H**). We hypothesize that the strong ability to transfect the OSCC lines is due to the isopropyl terminal branching at the end of the ionizable lipid tails, which we found to aid in mRNA endosomal escape (**Figure 2I**). To further probe potency, we reformulated E10i-494 with mCherry mRNA and performed a dose escalation study in all three cell lines, ranging from 0.05 ng to 250 ng per 250,000 cells. Differences were observed in all cell lines, with CAL-27 being the easiest to transfect as a 0.25 ng dose resulted in 50% cell transfection and 80% cells were transfected at a 5 ng dose (**Figure 2J–K**). Meanwhile, OECM-1 cells were the most difficult to transfect as a 25 ng dose only resulted in 60% transfection (**Figure 2L–M**). Lastly, FaDu cells had a medium transfectability, with 1 ng inducing 20% positivity while 5 ng resulted in 80% positivity (**Figure 2N–O**).

### Co-loading of p53 mRNA and CPX results in synergistic cell killing in vitro and tumor reduction in vivo

After identifying E10i-494 as the lead candidate, we reformulated the LNP with human p53 mRNA to assess whether the mRNA can induce OSCC killing in vitro. Upon comparing E10i-494 formulated with FLuc mRNA (FLuc-LNP) compared to human p53 mRNA (p53-LNP), we found that the p53 mRNA results in dose-dependent cellular toxicity in CAL-27 (**Figure 3A**), OECM-1 (**Figure 3B**), and FaDu (**Figure 3C**) cells. A further increase in p53-induced toxicity was observed at 48 h (**Supplementary Figure 2A–C**), indicating long-lasting cell killing. Interestingly, FLuc-LNP induced toxicity in CAL-27 cells, while in OECM-1 and FaDu cells, no FLuc-LNP toxicity was observed. This is likely a result of the cellular differences among the OSCC lines, as observed with the mCherry results, where some cell lines may be more sensitive to LNP delivery. When comparing E10i-494 encapsulating p53 mRNA against PTEN mRNA, another tumor suppressor gene commonly inactivated in OSCC, the p53-LNP induced 2–4-fold greater toxicity than with PTEN mRNA (PTEN-LNP) in all three OSCC lines (**Supplementary Figure 3**). With the efficacy of p53 mRNA demonstrated in vitro, we next evaluated CPX. Since CPX has not been evaluated in OSCC lines, we performed a dose response assay with free CPX and found that the free drug reduced viability to 50% at concentrations around 100 ng per 20,000 cells in all three cell lines (**Supplementary Figure 4A–C**). As CPX has low water solubility and high solubility in ethanol, we formulated four LNPs encapsulating CPX at 2, 10, 25, and 50 mole percent (mol%) with FLuc mRNA by dissolving CPX in the ethanol phase along with the other lipid excipients. High CPX incorporation within the LNP resulted in less mRNA encapsulation, with the 50 and 25 mol% CPX LNPs having 70% and 65% mRNA encapsulation, respectively, compared to ~80% encapsulation for the 10 mol%, 2 mol%, and CPX-free LNPs (**Supplementary Figure 5A**). Similarly, the 50 and 25 mol% CPX LNPs had smaller hydrodynamic diameters of ~70 nm compared to 80–90 nm for the other CPX and CPX-free LNPs (**Supplementary Figure 5B**). These alterations in LNP physical properties indicate that CPX is successfully being co-encapsulated with mRNA into the LNP.

The four CPX LNPs were then evaluated for their chemotherapeutic properties by incubating them in CAL-27, OECM-1, and FaDu cells at doses of 20 ng/20,000 cells. While no decrease in viability was observed at 24 h, at 48 h, the 50 and 25 mol% CPX LNPs resulted in viabilities of ~75%, a modest but noticeable decrease in viability (**Supplementary Figure 6A– C**). To determine the synergistic properties of CPX and p53, we reformulated E10i-494 with p53 mRNA containing either 25 mol% CPX (25% CPX/p53-LNP) or 50 mol% CPX (50% CPX/p53-LNP). Upon dosing in the three cell lines at 2, 5, and 20 ng per 20,000 cells, we observed improved cell killing in all three cell lines for the 50% CPX/p53-LNP compared to the p53-LNP, especially at the 5 ng dose, whereas 25% CPX/p53-LNP facilitated similar or higher viability compared to p53-LNP. The greatest disparities in viability were observed at the 5 ng. For CAL-27 cells at a 5 ng, the addition of 50 mol% CPX resulted in a decrease from 50% to 30% viability compared to p53-LNP, whereas for OECM-1 and FaDu cells, the viability decreased from 100% to 65% (**Figure 3D–F**). With the 50 mol% CPX LNP inducing the lowest viability, 50 mol% was incorporated for all LNPs containing CPX (CPX-LNP) as well as for formulations containing p53 mRNA and CPX (CPX/p53-LNP) throughout the rest of the studies.

**Figure 3.**
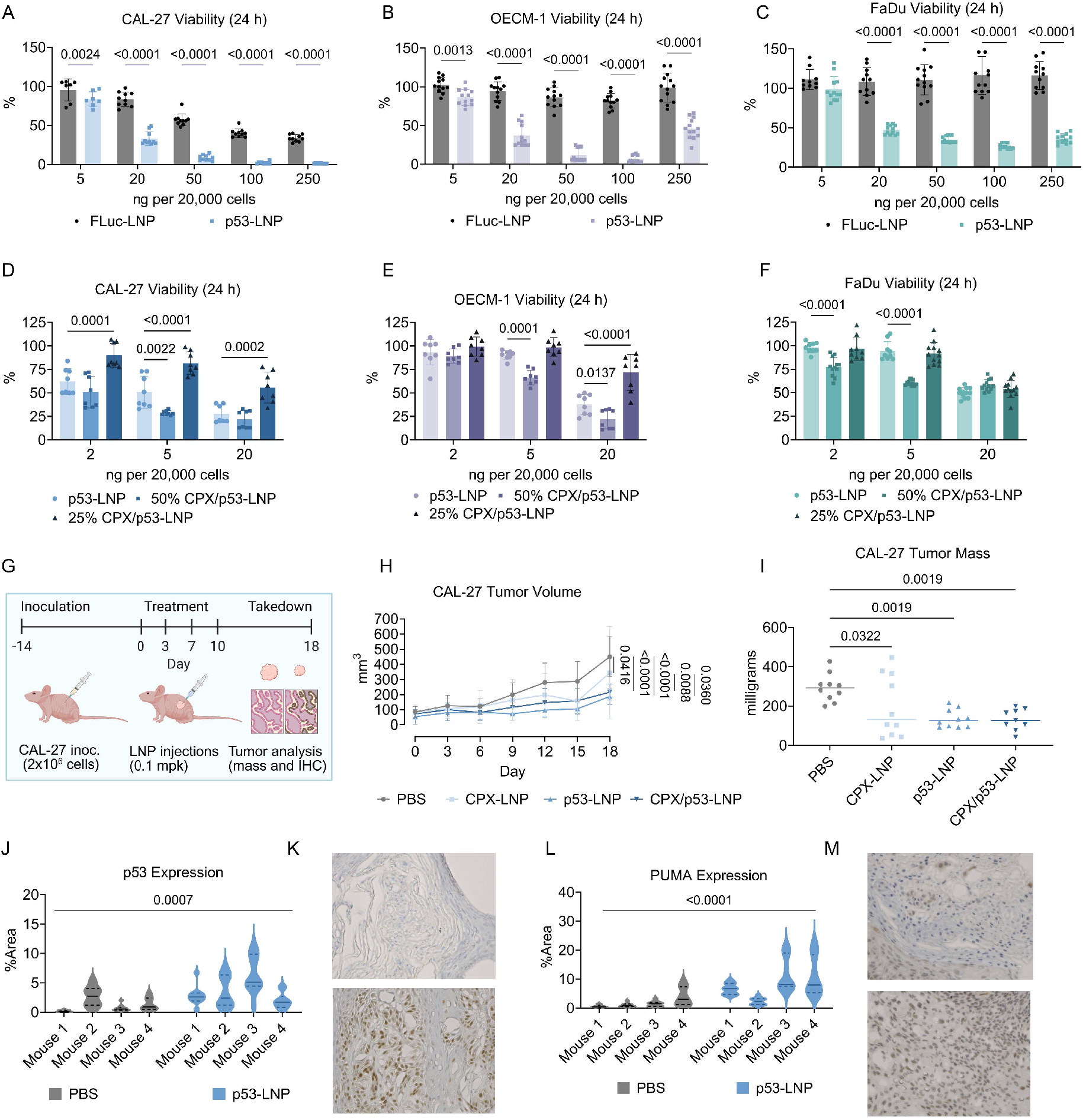
Cell killing properties of E10i-494 p53 mRNA and CPX co-therapy. **A–C**, E10i-494 was formulated with FLuc mRNA (FLuc-LNP) and separately with human p53 mRNA (p53-LNP). The two LNPs were incubated in **(A)** CAL-27, **(B)** OECM-1, and **(C)** FaDu cells at increasing dosages per 20,000 cells, and after 24 h, viability was measured. Viability is reported as mean ± SD of *n* = 8– 12. **D–F**, p53-LNP, E10i-494 reformulated with p53 mRNA and 50 mol% CPX (50% CPX/p53-LNP), and E10i-494 reformulated with 25 mol% CPX (25% CPX/p53-LNP) were incubated in **(D)** CAL-27, **(E)** OECM-1, and **(F)** FaDu cells at 2, 5, and 20 ng of mRNA per 20,000 cells, and after 24 h, viability was measured. Viability is reported as mean ± SD of *n* = 8 for CAL-27 and OECM-1 cells and *n* = 9 for FaDu cells. **G**, Nu/J mice were inoculated with two million CAL-27 cells on each flank, and after fourteen days, LNPs were administered intratumorally at an mRNA dose of 0.1 mpk four times over ten days in each tumor. **H–N**, Throughout the study, **(H)** tumor size was measured, and after eighteen days, the mice were sacrificed and **(I)** tumors were weighed. The PBS and p53-LNP groups were further analyzed for protein expression, examining **(J)** p53 protein from the corresponding **(K)** IHC images as well as **(L)** PUMA protein from the corresponding **(M)** IHC images. Tumor volume and mass are reported as mean ± SD of n = 9 tumors from *n* = 5 mice. Protein expression is measured from *n* = 10 images with a solid representing the median and dashed lines representing quartiles. Representative IHC images are shown for each group. **A–C**, Two-tailed multiple unpaired t test with *post hoc* Holm–Šídák correction for multiple comparisons was used to compare FLuc-LNP to p53-LNP. **D–F**,**H**,**J**,**L**, Two-way ANOVA with *post hoc* Holm–Šídák correction for multiple comparisons was used to compare **(D–F)** p53-LNP viability across LNP types for each dose, **(H)** tumor volume at each time point across groups, and **(J)** p53 and **(L)** PUMA %area between PBS and p53-LNP groups. **I**, One-way ANOVA with *post hoc* Holm–Šídák correction for multiple comparisons was used to compare tumor mass to PBS.

With the chemotherapeutic properties of p53 mRNA and CPX demonstrated in vitro, we moved to a proof-of-concept in vivo study. Two million CAL-27 cells were inoculated in Nu/J mice on each flank, and after two weeks, PBS, CPX-LNP, p53-LNP, and CPX/p53-LNP were intratumorally administered at 0.1 mg of mRNA per kg of body weight (mg/kg) four times over two weeks on each tumor (**Figure 3G**). Tumor volumes were measured throughout the course of the experiment and after a total of eighteen days from the first injection, the mice were sacrificed, tumor mass was measured, and some of the tumors were processed using immunohistochemistry (IHC). While the CPX-LNP induced a modest decrease in tumor volume and mass, the p53-LNP and CPX/p53-LNP significantly delayed tumor growth by one-third of the PBS burden, demonstrating that for CAL-27 xenograft tumors, p53 mRNA and not CPX, is sufficient to diminish tumor burden (**Figure 3H–I**). Additionally, no weight loss was observed (**Supplementary Figure 7**). IHC analysis of tumor tissues from PBS control and p53-LNP-treated groups revealed that the latter had increased expression of p53 and its downstream target, PUMA, demonstrating that pro-apoptotic pathways were activated because of the therapy (**Figure 3J–M**).

### p53 mRNA and CPX activate caspases in cancer cells and repolarize tumor-associate macrophages

To probe the mechanism of tumor cell death and investigate whether the therapies can modulate the tumor microenvironment, we performed a series of mechanistic studies. First, we measured caspase activity in tumor cells. Caspase enzymes are cysteine proteases that facilitate apoptosis, which can be triggered extracellularly and intracellularly. Many chemotherapeutics, once delivered into the cell, mediate intracellular caspase-based apoptosis, where mitochondria release cytochrome C that induces the formation of an apoptosome, a complex protein complex that recruits and activates caspase-9, an initiator caspase^26^. Once caspase-9 is engaged, a caspase activation cascade occurs, leading to the activation of executioner caspases, including caspase-3 and caspase-7, and consequently, the cell death. To explore the combinatory effect of p53 and CPX on caspase activity, FLuc-LNP, p53-LNP, CPX-LNP, and CPX/p53-LNP were dosed in CAL-27, OECM-1, and FaDu cells at 5, 20, and 50 ng per 20,000 cells. After 24 h, a fluorescence-based caspase-3/7 assay was conducted, which measures caspase-3/7 activity. Normalizing to untreated wells, we found that in CAL-27 cells, p53-LNP increases caspase-3/7 activity in each dose, but for the 50 ng dose, the CPX/p53-LNP group had 10% higher fluorescence (**Figure 4A**), indicating CPX increases caspase-3/7 activity in CAL-27 cells. In contrast, CPX did not enhance caspase-3/7 activity in OECM-1 and FaDu cells, as the p53-LNP and CPX/p53-LNP groups maintained the same fluorescence at all doses (**Figure 4B–C**). Out of the three cell lines, FaDu cells produced the most caspase-3/7 upon LNP treatment, as the p53-LNP and CPX/p53-LNP groups produced ~2.5-fold higher fluorescence than baseline, compared to 1.5-fold and 1.25-fold in CAL-27 and OECM-1 cells, respectively. We then examined caspase-9 activity utilizing a luminescence-based reporter system. As the 50 ng dose induced the strongest caspase-3/7 activity, we dosed the three cell lines for the caspase-9 studies at 50 ng. After 24 h, we observed an opposite trend from the previous study. Here, CPX/p53-LNP enhanced caspase-9 activity by 2-fold and 1.5-fold compared to p53-LNP, whereas there were no differences in activity for the two LNPs in CAL-27 cells (**Figure 4D**). Similar to the caspase-3/7 assays, treatment in FaDu cells induced the largest amount of caspase-9 compared to the other cell lines. Thus, CPX enhances p53-mediated caspase activity, but the specific caspase depends on the cell line. When examining other potentially related pathways, we found that none of the LNPs induce meaningful increases in mitochondrial permeability (**Supplementary Figure 8**) nor reactive oxygen species (**Supplementary Figure 9**).

**Figure 4.**
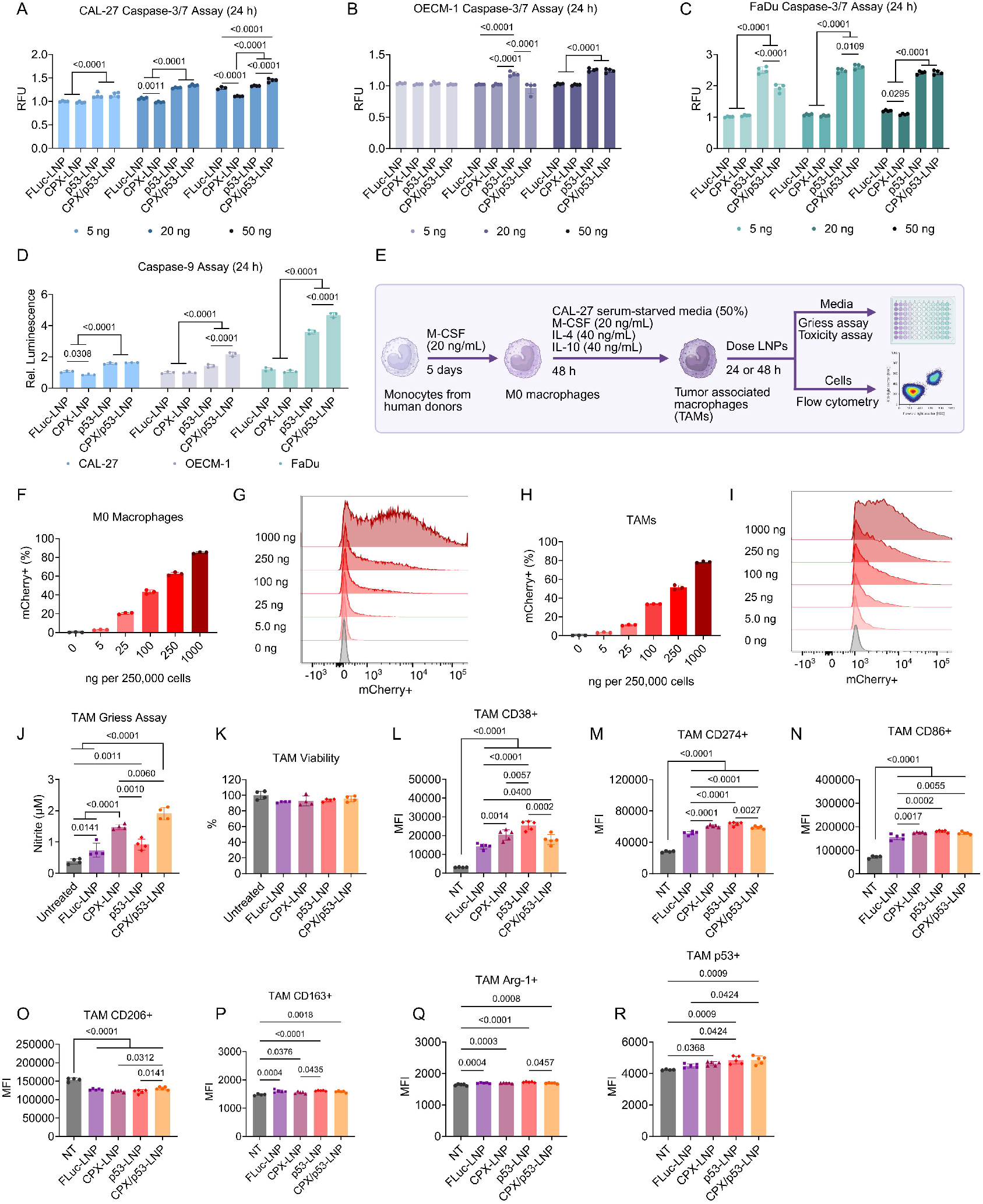
Mechanistic analysis of cell killing and macrophage polarization for CPX/p53-LNP platform. **A–C**, E10i-494 LNPs containing FLuc mRNA (FLuc-LNP), 50 mol% CPX (CPX-LNP), human p53 mRNA (p53-LNP), and 50 mol% CPX and human p53 (CPX/p53-LNP) were dosed in **(A)** CAL-27, **(B)** OECM-1, and **(C)** FaDu cells at 5, 20, and 50 ng per 20,000 cells. After 24 h, a fluorescence-based caspase-3/7 assay was performed on the cell supernatant, where fluorescence was calculated relative to untreated cells. Relative fluorescent units (RFU) are reported as mean ± SD of *n* = 4. **D**, All four LNPs were incubated in CAL-27, OECM-1, and FaDu cells at a dose of 50 ng per 20,000 cells. After 24 h, a luminescence-based caspase-9 assay was performed on cell supernatant, where luminescence was calculated relative to untreated cells. Relative luminescence is reported as mean ± SD of *n* = 3. **E**, Schematic of primary human monocyte polarization to M0 macrophages and TAMs using a combination of cytokines and serum-depleted media from CAL-27 cells. **F–I**, E10i-494 encapsulating mCherry mRNA was incubated with M0 macrophages and TAMs at increasing doses per 250,000 cells. After 24 h, flow cytometry was utilized to determine **(F)** mCherry+ M0 macrophages, **(G)** M0 macrophage fluorescence intensity, **(H)** mCherry+ TAMs, and **(I)** TAM fluorescence intensity. mCherry positivity is reported as mean ± SD of *n* = 3. Representative data are shown for fluorescence intensity plots. **J–R**, All four LNPs were incubated in TAMs at a dose of 250 ng per 250,000 cells, and **(J)** after 24 h, a Griess assay was performed to measure nitrite production and **(K)** a viability assay was employed to determine TAM viability, while after 48 h, flow cytometry was used to examine **(L)** CD38 MFI, **(M)** CD274 MFI, **(N)** CD86 MFI, **(O)** CD206 MFI, **(P)** CD163 MFI, **(Q)** Arg-1 MFI, and **(R)** p53 MFI. Nitric oxide concentration and cellular viability are reported as mean ± SD of *n* = 4 while protein MFI is reported as mean ± SD of *n* = 5. **A–D**, Two-way ANOVA with *post hoc* Holm– Šídák correction for multiple comparisons was used to compare **(A–C)** RFU across LNP types for each dose and **(D)** relative luminescence across LNP types for each cell line. **J–R**, One-way ANOVA with *post hoc* Holm–Šídák correction for multiple comparisons was used to compare **(J)** nitrite concentration, **(K)** cellular viability, and **(L–R)** MFI across each LNP type.

To investigate the effect of the therapies on TIME, we examined macrophage polarization, focusing on tumor-associated macrophages (TAMs) as the dominant innate immune cells in OSCC tumors, which are associated with poorer outcomes^27^. We first obtained monocytes from healthy human donors and incubated them with macrophage colony-stimulate factor (M-CSF) for five days to form M0 macrophages (**Figure 4E**). To avoid unintentional polarization, we cultured the cells in human serum, rather than bovine serum, to mimic more realistic conditions. After converting to M0 macrophages, we further cultured the cells with IL-4, IL-10, and conditioned medium from serum-starved CAL-27 cells for 48 h to induce OSCC TAMs. To determine whether the E10i-494 LNP, which was optimized for OSCC lines, can transfect macrophages, we reformulated the LNP with mCherry mRNA and performed a dose escalation study in both M0 macrophages and TAMs. Although more difficult to transfect, mCherry expression was observed in a dose-dependent manner, where 80% of M0 macrophages and 75% of TAMs were mCherry positive at a dose of 1,000 ng per 250,000 cells after 24 h (**Figure 4F–I**). We then determined if the LNPs can promote nitric oxide formation, which is a pro-inflammatory molecule characteristic of M1-like macrophages, but is produced in minute concentrations in the TME^28^. TAMs were dosed at 250 ng per 250,000 cells with FLuc-LNP, CPX-LNP, p53-LNP, and CPX/p53-LNP for 24 h. Afterwards, we performed a Griess assay, which examines nitrite, as nitric oxide is rapidly converted to nitrite in cell media. Using nitrite concentration as a proxy for nitric oxide, we observed that CPX-LNP and CPX/p53-LNP increased nitric oxide production in TAMs by 1.5-fold and 2-fold, respectively, compared to p53-LNP, and 3-fold and 4-fold compared to FLuc-LNP (**Figure 4J**). This suggests that CPX is the major driver of nitric oxide production, although p53, when combined with CPX, can further increase production. In addition, no obvious changes in cell viability were observed after dosing in TAMs, suggesting that CPX-LNP and p53-LNP had no effects on cell viability of macrophages (**Figure 4K**). A slight increase in nitric oxide production was observed in FLuc-LNP compared to untreated cells, demonstrating that the innate inflammatory nature of LNPs can also induce nitric oxide formation.

To probe how the LNPs modulate TAM phenotype, we repeated the treatment regimen and examined the cells by flow cytometry after 48 h. We found that both the LNP itself and the encapsulated p53 mRNA contribute separately to repolarizing TAMs into a more M1-like phenotype. The median fluorescent intensity (MFI) of M1-like markers such as CD38 and CD274 was increased by CPX-LNP, p53-LNP, and CPX/p53-LNP, but most strongly by p53-LNP (**Figure 4L–M**). The M1-like marker CD86 was elevated for all groups, including FLuc-LNP, indicating that the inflammatory nature of the LNP was the driving factor for CD86 upregulation (**Figure 4N**). The MFI of the M2-like marker CD206 was also lowered by all LNPs, whereas for CD163 and arginase-1 (Arg-1), only minor differences in MFI were observed (**Figure 4O–Q**). Lastly, both p53-LNP and CPX/p53-LNP facilitated greater p53 MFI in TAMs, due to the delivery of p53 mRNA, while CPX-LNP slightly increased p53 MFI (**Figure 4R**). The flow results and Griess assay describe a complex interaction between the LNP, CPX, and p53 mRNA, where each is contributing to the repolarization of TAMs to a more M1-like phenotype. Although further studies are needed to better define the mechanism, these preliminary results suggest that CPX can increase the secretion of pro-inflammatory nitric oxide, while p53 mRNA and the LNP vehicle itself alter TAM phenotypes.

### CPX is the major driver of tumor reduction in p53-resistant MOC-2 syngeneic tumors

Having demonstrated the potency of the platform in human OSCC, we then evaluated whether this co-therapy approach could improve outcomes in a syngeneic murine tumor model formed by MOC-2 cells, an aggressive mouse OSCC line. We first reformulated E10i-494 with mCherry mRNA and incubated the LNP in MOC-2 cells at increasing doses to evaluate the transfection efficiency. We found that at 25 ng per 250,000 cells, E10i-494 transfected almost 80% of MOC-2 cells, which is on par with the human OSCC lines (**Figure 5A–B**). However, when assessing the potency of CPX-and p53-loaded LNPs in MOC-2 cells, the latter using a mouse p53 template, after 24 h, 250 ng was required to elicit cell death – a ten-fold greater dose than that required for the human OSCC lines (**Figure 5C**). After 48 h, there were further increases in toxicity at the 250 ng dose, with some increase at the 100 ng dose (**Supplementary Figure 10**). No significant changes in caspase-9 production were observed at doses of 5, 20, and 50 ng per 20,000 cells for all LNPs (**Figure 5D**), but an increase in caspase-3/7 activity was observed upon treatment with the highest dose of p53-LNP and CPX/p53-LNP, indicating functional p53 mRNA (**Figure 5E**). We then examined whether our therapeutic system could extend survival in C57BL/6J mice inoculated with MOC-2 cells. LNPs were administered at a dose of 1.0 mpk four times over ten days, and mice were monitored for up to forty days from the initial injection (**Figure 5F**). No loss in mouse weight was observed during the LNP administration period (**Supplementary Figure 11**). After twenty-two days from the first injection, the CPX-LNP and CPX/p53-LNP groups had lower tumor volumes than the PBS, FLuc-LNP, and p53-LNP groups, with the co-therapy inducing the lowest tumor volume (**Figure 5G**). The CPX-LNP and CPX/p53-LNP groups also maintained a flatter tumor growth curve during the forty-day experiment compared to the other groups (**Figure 5H–L**). p53-LNP only slightly prolonged survival, but CPX-LNP and CPX/p53-LNP increased survival by over a week, suggesting that CPX can enhance anti-tumor effect and prolong survival in mice burdened with tumors that respond poorly to p53 therapy (**Figure 5M**). To determine the mechanism underlying the reduction in tumor burden in this syngeneic model, we subcutaneously inoculated MOC-2 cells into new C57BL/6J mice (**Figure 5N**). The four LNPs were injected intratumorally into MOC-2 tumors four times over a ten-day period at a dose of 0.5 mpk, and after sixteen days from the initial injection, tumors were harvested for flow cytometry and IHC. Flow cytometry revealed that the LNPs induced a shift to a more M1-like phenotype, specifically for a unique class of CD11b+, F4/80-macrophages within the tumors, as there were decreases in Arg-1 and CD206 expression combined with increases in CD86 and CD274 expression (**Figure 5O–S**). As expected, IHC staining showed a sharp increase in p53 expression in tumor tissues treated with p53-LNP and CPX/p53-LNP (**Figure 5T**).

**Figure 5.**
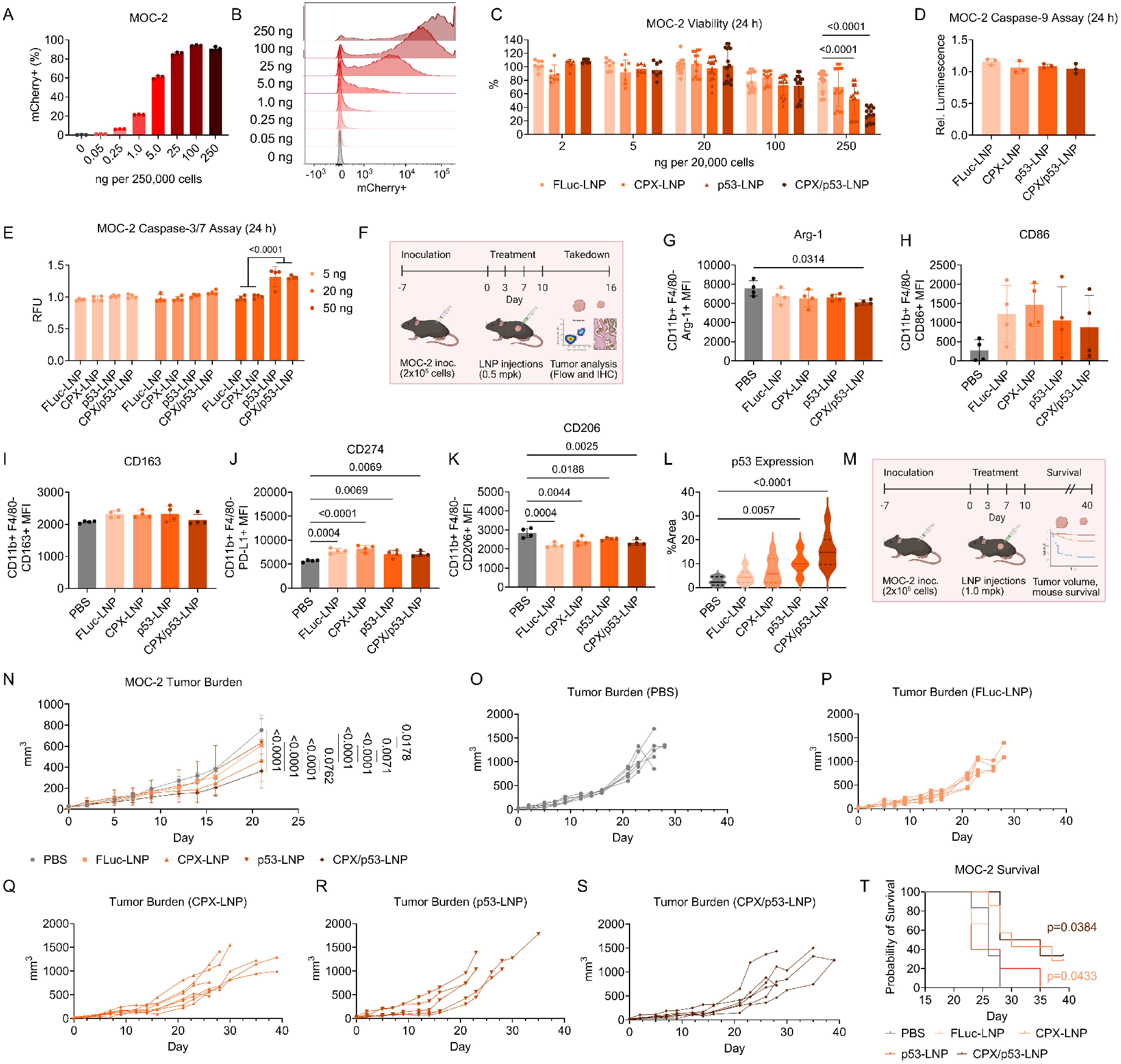
Evaluation of LNP co-therapy in a syngeneic model. **A–B**, E10i-494 encapsulating mCherry mRNA was incubated with MOC-2 cells at increasing doses per 250,000 cells. After 24 h, flow cytometry was utilized to determine **(A)** mCherry+ MOC-2 cells and **(B)** MOC-2 fluorescence intensity. mCherry positivity is reported as mean ± SD of *n* = 3. Representative data are used for the fluorescence intensity plots. **C**, E10i-494 LNPs containing FLuc mRNA (FLuc-LNP), 50 mol% CPX (CPX-LNP), mouse p53 mRNA (p53-LNP), and 50 mol% CPX and mouse p53 (CPX/p53-LNP) were incubated in MOC-2 cells at increases doses per 20,000 cells, and after 24 h, viability was measured. Cellular viability is reported as mean ± SD of *n* = 8 for 2 ng and 5 ng doses and *n* = 12 for all other doses. **D**, All four LNPs were incubated in CAL-27, OECM-1, and FaDu cells at a dose of 50 ng per 20,000 cells. After 24 h, a luminescence-based caspase-9 assay was performed on cell supernatant, where luminescence was calculated relative to untreated cells. Relative luminescence is reported as mean ± SD of *n* = 3. **E**, All four LNPs were incubated in CAL-27, OECM-1, and FaDu cells at doses of 5, 20, and 50 ng per 20,000 cells. After 24 h, a fluorescence-based caspase-3/7 assay was performed on the cell supernatant, where fluorescence was calculated relative to untreated cells. Relative fluorescent units (RFU) are reported as mean ± SD of *n* = 4. **F**, C57BL/6J mice were inoculated with 20,000 MOC-2 cells on the right flank, and after seven days, LNPs were administered intratumorally at an mRNA dose of 1.0 mpk four times over ten days. **G–L**, Seven days after the initial inoculation, tumor volumes were measured **(G)** comparatively for up to twenty-two days as well as individually for the **(H)** PBS, **(I)** FLuc-LNP, **(J)** CPX-LNP, **(K)** p53-LNP, and **(L)** CPX/p53-LNP groups. Tumor burden is reported as mean ± SD of *n* = 6 for PBS and *n* = 5 for all other groups. **M**, Kaplan-Meier survival curves for all five groups. **N**, C57BL/6J mice were inoculated with 20,000 MOC-2 cells on the right flank, and after seven days, LNPs were administered intratumorally at an mRNA dose of 0.5 mpk four times over ten days. **O–S**, After sixteen days from the initial injection, the mice were sacrificed, and the tumors were harvested and analyzed by flow cytometry to examine **(O)** Arg-1 MFI, **(P)** CD86 MFI, **(Q)** CD163 MFI, **(R)** CD274 MFI, and **(S)** CD206 MFI. MFI is reported as mean ± SD of *n* = 4. **T**, Additionally, tumors were characterized by IHC for p53 expression. Protein expression is measured from *n* = 8 images with a solid line representing the median and dashed lines representing quartiles. **C**,**E**,**G**, Two-way ANOVA with *post hoc* Holm–Šídák correction for multiple comparisons was used to compare **(C)** cellular viability across LNP types for each dose, **(E)** RFU activity across LNP types for each dose, and **(G)** tumor volume across LNP types for time point. **D**,**O–T**, One-way ANOVA with *post hoc* Holm–Šídák correction for multiple comparisons was used to compare **(D)** relative luminescence, **(O–S)** MFI, and **(T)** %area across each LNP type. **M**, Kaplan-Meier estimator with post hoc Holm-Šídák correction for multiple comparisons was used to compare survival curves to PBS.

## Discussion

OSCC is a debilitating disease due to the importance and multifunctionality of the oral cavity. Mainline treatments including surgery and chemoradiation can halt disease progression but come at the cost of further damage to the oral cavity, where in most cases, the tongue is either partially or completely resected. Moreover, many patients have also not benefited from current state-of-the-art immunotherapies, including Cetuximab and checkpoint blockade inhibitors due to a lack of response. While other cellular and immunotherapies have facilitated remission for different cancers, OSCC has largely been left out of testing. Thus, we aimed to develop a co-therapy platform based on the biology of OSCC development, by targeting both the inactivation of TSGs and the immunosuppressive TIME. A hallmark of OSCC is that its progression relies more on the inactivation and mutation of TSGs, rather than the overexpression of multiple oncogenes. This allows it to be well-suited for mRNA-based approaches that can facilitate the expression and functional activation of TSGs in OSCC.

LNPs are the primary vehicle for mRNA delivery in the clinic; however, the complex structure of LNPs also allows for encapsulation of therapeutics other than nucleic acids^29^. For hydrophilic compounds, intermolecular interactions, such as electrostatic interactions, with the LNP components must be leveraged to achieve successful encapsulation, as otherwise, the compound will be mainly located outside of the LNP in the surrounding buffer because of favorable entropic and enthalpic conditions. Similarly, therapeutics that are too hydrophobic may not dissolve in ethanol and thus are unable to be incorporated into the LNP. CPX is uniquely situated to be encapsulated due to its combination of low water solubility and high solubility in ethanol, allowing it to be co-dissolved in the ethanolic phase along with the other lipid components. There are likely many other drug candidates that have favorable solubility properties and can also be formulated with nucleic acids for potent co-therapy. This pool of small molecule drugs can be expanded by altering the aqueous and organic phases prior to LNP formulation.

Our approach not only investigated delivery of two therapeutics but also examined the role of simultaneous uptake in cancer cells and macrophages. Many cancer cells readily take up LNPs due to their high metabolism and dependence on cholesterol for membrane biogenesis; however, macrophages are also a repository for LNPs due to their phagocytic nature^30^. As TAMs are one of the dominant immune cells in the oral TME, devising nanocarriers to both kill cancer cells and reshape the TIME would greatly increase the potency of the drug delivery platform. While p53 has been demonstrated to repolarize M2-like macrophages to M1-like macrophages, our results suggest, for the first time, that p53 can repolarize TAMs within the oral TIME of OSCC^15^. This is significant since previous p53-based therapies have focused solely on targeting cancer cells, meaning that possible benefits from targeting immune cells have been overlooked. Similar to the growing evidence that traditional chemotherapeutics, including doxorubicin, activate the immune system, we demonstrate that p53 likewise possesses a similar chemoimmunotherapeutic potential, although more studies are needed to further elucidate this mechanism^31^. Secondly, our results demonstrated that CPX can polarize TAMs into M1-like phenotypes characteristic of increased nitric oxide production. Even though CPX has shown immunomodulatory effects in certain settings, including boosting T cell infiltration in a syngeneic mouse 4T1 breast tumor model^32^ and suppressing osteoclast formation in an ovariectomy-Induced osteoporosis model in mice^33^, little is known about its role in macrophage polarization. A recent study showed that blocking spermidine-mediated hypusination of the eukaryotic translation initiation factor 5A (eIF5A) inhibited OSCC-induced M2-like TAMs^34^. Several studies demonstrate that CPX exerts its anti-tumor effects by inhibiting the activity of deoxyhypusine hydroxylase (DOHH), one of the key enzymes responsible for eIF5A hypusination^35,36^. Further studies are warranted to explore whether CPX skews OSSC-induced M2-like TAMs into a M1-like phenotype by blocking DOHH-mediated eIF5A hypusination.

Although the cell killing potential of p53 is well-established, we found that CPX alone encapsulated in an LNP is a poor chemotherapeutic, yet in combination with p53 mRNA, CPX enhances caspase activity. More interestingly, the specific caspase affected by CPX depended on the OSCC line derived from different anatomical regions of the oral cavity, suggesting that each OSCC line has a unique apoptosis pathway due to their intrinsic heterogeneity. These results emphasize the need for testing multiple heterogeneous cell lines for OSCC, as well as other types of cancer cell lines, to properly gauge how the therapeutic platform will respond to phenotype differences. This is particularly important for translating a product to human OSCC, which is highly heterogeneous. The patient-to-patient variability is a significant reason that previous p53-based therapies have failed in the clinic, as most tumors are p53 deficient but not all cancer cells in a tumor may be p53-depleted, while others may have mutated p53 that overrides wild type p53^37^. To account for this, we tested our platform in MOC-2 cells, a murine OSCC line, which we found to be resistant to mouse p53 mRNA. Despite this tolerance, the addition of CPX was able to reduce tumor burden and extend survival in a MOC-2 syngeneic mouse model due to its chemotherapeutic and immunomodulatory behavior. Thus, this platform has the potential to treat OSCC broadly, even in cases where one of the therapies is ineffective.

In summary, we have identified an optimal LNP for transfection in OSCC through a multi-tiered approach by screening ionizable lipids with unique architecture for potent transfection and low liver delivery. We then leveraged the hydrophilic and hydrophobic domains in LNPs to co-deliver p53 mRNA and the small molecule CPX. The therapies display synergistic anti-tumor effects by accelerating caspase activity. Additionally, the LNP can transfect OSCC TAMs, where the combination of p53, CPX, and the LNP itself induces polarization of OSCC-induced TAMs to an M1-like phenotype. When evaluated in p53-susceptible and p53-resistent models, we found that the co-delivery strategy can reduce tumor burden in both p53-sensitive and resistant OSCC models.

## Methods

All animal use was in accordance with the guidelines and approval from the University of Pennsylvania’s Institutional Animal Care and Use Committee (IACUC; protocols 806540 and 804466).

### Materials

All non-synthesized lipid excipients were purchased from Avanti Polar Lipids (Alabaster, AL, USA). Firefly luciferase mRNA with 5-methoxyuridine substitutions and mCherry mRNA with N1-methylpseudouridine substitutions were purchased from TriLink Biotechnologies (San Diego, CA, USA). 1,2-epoxyoctane, 1,2-epoxydecane, 1,2-epoxydodecane, and 1,2-epoxytetradecane were purchased from TCI (Montgomeryville, PA, USA); Triton X-100 was purchased from Alfa Aesar (Haverhill, MA, USA). 2-{2-[4-(2-{[2-(2-aminoethoxy)ethyl]amino}ethyl)piperazin-1-yl]ethoxy}ethan-1-amine (494) was purchased from Enamine (Kyiv, Ukraine). Ciclopirox and all other chemical reagents were purchased from MilliporeSigma (St. Louis, MO, USA).

### Synthesis

Flash chromatography was performed on a Teledyne ISCO (Lincoln, NE, USA) CombiFlash NextGen 300+ equipped with evaporative light scattering detection (ELSD) using RediSep Gold silica gel disposable flash columns (Teledyne ISCO). Solvent evaporation was performed using a Büchi (New Castle, DE, USA) Rotavapor R-300 System Professional. ^1^H and ^13^C NMR spectra were acquired in chloroform-d using an Avance Neo 400 MHz spectrometer (Bruker, Billerica, MA, USA). NMR spectra were analyzed on MestReNova 14.3.0 (Mestrelab, Santiago de Compostela, Spain). Nominal mass accuracy LC-MS data were obtained by use of a Waters (Milford, MA, USA) Acquity UPLC system equipped with a Waters TUV detector (254 nm) and a Waters SQD single quadrupole mass analyzer with electrospray ionization. Samples were prepared in 200 proof ethanol and injected onto an Acquity UPLC BEH C8 1.7 µm, 2.1 × 50 mm column with a 2 min wash followed by a gradient mobile phase from 50% water (1% trifluoroacetic acid) and 50% acetonitrile (1% trifluoroacetic acid) to 100% acetonitrile (1% trifluoroacetic acid) over 8 min. LC-MS spectra were analyzed on MestReNova 14.3.0.

To a 3 mL vial (Thermo Fisher Scientific) was added the 494 polyamine core (1.0 equiv.), the corresponding epoxide (7.0 equiv.), and ethanol (0.3 mL). The reaction stirred at 80 °C for 48 h. Afterwards, the crude product was diluted with dichloromethane (0.7 m) and purified via flash chromatography assisted by CombiFlash using a liquid injection into a 4 g RediSep Gold silica disposable flash column (Teledyne Isco). The mobile phase consisted of a gradient of 95% dichloromethane and 5% Ultra solution (75% dichloromethane, 22% methanol, and 3% aqueous ammonium hydroxide) to 80% dichloromethane and 20% Ultra solution over 35 min using a flow rate of 7 mL/min. The products were isolated as yellow to clear viscous oils. See Padilla *et al*.^23^ for the full synthetic pathway, specific conditions, and spectra.

### mRNA production

Codon-optimized human p53, human PTEN, and mouse p53 sequences were cloned into a plasmid with optimized 3′ and 5′ untranslated regions (UTRs) and containing a 101 polyA tail. mRNA was produced via in vitro transcription of the plasmid using 1-methylpseudouridine-modified nucleosides and co-transcriptionally capped with CleanCap technology (TriLink). The mRNA was purified with cellulose to remove double-stranded RNAs, and purified mRNAs were precipitated in ethanol and were re-suspended in nuclease-free water. Electrophoresis, dot blot, and endotoxin assays were performed for quality control. All mRNAs were stored in a −80 °C freezer until use.

### Lipid nanoparticle formulation

Cholesterol, 18:1 Δ9-cis phosphoethanolamine (DOPE), 14:0 PEG2000 phosphoethanolamine (C14-PEG_2000_), and an ionizable lipid were dissolved in ethanol at a molar ratio of 46.5:16:2.5:35, respectively. CPX was added to the ethanol phase for the LNPs containing CPX. The SM-102 formulation was prepared using cholesterol, 1,2-distearoyl-sn-glycero-3-phosphocholine (DSPC), and 1,2-dimyristoyl-rac-glycero-3-methoxypolyethylene glycol-2000 (DMG-PEG_2000_), and the SM-102 ionizable lipid at a molar ratio of 38.5:10:1.5:50, respectively. The ALC-0315 formulation was prepared by combining cholesterol, DPSC, ALC-0159, and the ALC-0315 ionizable lipid at a molar ratio of 42.7:9.4:1.6:46.3, respectively. The aqueous phase was prepared by dissolving the corresponding mRNA in 10 mM citrate buffer at pH 3 (Teknova, Hollister, CA, USA). The volume ratio of the ethanol phase to aqueous phase was 1:3 with a 10:1 weight ratio of the corresponding ionizable lipid and mRNA. The citrate and ethanol phases were loaded into glass syringes (Hamilton Company, Reno, NV, USA) and connected to a Pump 33 DDS syringe pump (Harvard Apparatus, Holliston, MA, USA) attached to a microfluidic device with a staggered herringbone micromixer design. Microfluidic devices were fabricated in polydimethylsiloxane utilizing standard soft lithographic procedures^24^. A two-step exposure process was used to create the SU-8 master with positive channel features on a silicon wafer, where each mixing channel is 4 cm in length. The syringes were injected into the microfluidic device using flow rates of 1.8 mL/min and 0.6 mL/min for the aqueous and ethanol phases, respectively.

After formulation, LNPs dialyzed against 1X PBS at pH 7.4 in 0.5 mL or 3 mL 20 kDa molecular weight cutoff (MWCO) cassettes (Thermo Fisher Scientific) for at least 2 h at room temperature. After dialysis, the LNPs were passed through a 0.22 µm syringe filter (Genesee Scientific, El Cajon, CA, USA) and stored at 4 °C until further use. When necessary, LNPs were concentrated using Amicon 50k regenerated cellulose centrifugal filters (Millipore Sigma) pre-rinsed with 1X PBS, centrifuging at 700 g until the desired volume was reached.

### Lipid nanoparticle characterization

Relative encapsulation efficiency and encapsulated mRNA concentration were measured by a Quant-iT RiboGreen assay (Thermo Fisher Scientific). Each sample was diluted 100-fold in either 1X tris-EDTA (TE) buffer or 1X TE buffer containing 1% (v/v) Triton X-100. The Triton X-100 samples were mixed thoroughly and allowed to incubate for 5 min to achieve complete lysis of the samples. A standard curve was generated by diluting the corresponding mRNA used during formulation to concentrations ranging from 2.00 µg/mL to 31.3 ng/mL in 1X TE buffer. LNPs in TE buffer and LNPs in Triton X-100 were plated in quadruplicate, while the mRNA standards were plated in duplicate in black 96-well plates (Greiner, Kremsmünster, Austria). Afterwards, the RiboGreen fluorescent detection reagent was added per the manufacturer’s instructions. The plate was then wrapped in aluminum foil and shook on a plate shaker at 200 rpm for 5 min. Afterwards, fluorescence intensity was analyzed on an Infinite 200 Pro plate reader (Tecan, Morrisville, NC) using the Tecan i-control 3.9.1.0 software at an excitation wavelength of 485 nm and an emission wavelength of 525 nm. RNA concentration was determined via a standard curve estimated from a univariate least squares linear regression (LSLR). Relative encapsulation efficiency was calculated as 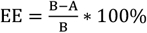, where A is the measured unencapsulated mRNA in TE buffer and B is the measured mRNA content after LNP lysis in Triton X-100. The encapsulated mRNA concentration was calculated by [conc] = B − A. Encapsulation efficiencies are reported as mean ± standard deviation.

The hydrodynamic diameters and polydispersity indexes (PDIs) of the LNPs were measured using a DynaPro Plate Reader III (Wyatt Technology, Santa Barbara, CA). The LNPs were diluted 10-fold in 1X PBS, and 30 µL were loaded onto a 384-well Aurora plate (Wyatt). The plate was centrifuged for 5 min at 300 g before loading onto the plate reader. Hydrodynamic diameters are reported as intensity-weighted averages with n = 3 measurements. Data is expressed as mean ± standard deviation, where the standard deviation was calculated by, 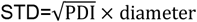. PDI is reported as mean ± standard deviation of the measurements. Data were analyzed using Dynamics 8.1.2.144 (Wyatt Technology).

### In vitro culturing

CAL-27 (CRL-2095) and FaDu (HTB-43) lines were purchased from ATCC (Manassas, VA, USA), MOC-2 (EWL002-FP) from Kerafast (Newark, CA, USA), and OECM-1 (SCC180) lines were purchased from MilliporeSigma. CAL-27, OECM-1, and FaDu cells were cultured in tissue-treated T75 flasks (Thermo Fisher Scientific) with Dulbecco’s modified Eagle medium with *L*-glutamine (DMEM; Gibco, Dublin, Ireland) supplemented with 10% fetal bovine serum (FBS; Gibco) and 1% penicillin-streptomycin (P/S; Gibco). MOC-2 cells were cultured in tissue-treated T75 flasks 2:1 IMDM (Thermo Fisher Scientific) and Hams Nutrient Mixture (Thermo Fisher Scientific) supplemented with 10% FBS, 1% P/S, 5 mg/L insulin (MilliporeSigma), 40 µg/L hydrocortisone (MilliporeSigma), and 5 µg/L epidermal growth factor (EGF; MilliporeSigma).

Primary human monocytes were obtained from healthy donors via the Perelman School of Medicine Human Immunology Core. The cells were incubated in non-tissue-treated T75 flasks containing Roswell Park Memorial Institute (RPMI) 1640 media (Gibco) supplemented with 10% human serum (MilliporeSigma), 1% P/S, and 20 ng/mL recombinant human macrophage colony-stimulating factor (M-CSF; PeproTech, Cranbury, NJ, USA) for five days at a maximum cell density of 1 million cells per mL to obtain M0 macrophages. The media was replaced every two or three days. To polarize M0 macrophages to TAMs, the M0 macrophages were cultured in 1:1 CAL-27 tumor conditioned media and RPMI supplemented with 10% human, 1% P/S, 20 ng/mL M-CSF, 40 ng/mL human recombinant IL-4 (PeproTech), and 40 ng/mL human recombinant IL-2 (PeproTech) for 48 h. CAL-27 tumor conditioned media was generated by plating 10 million CAL-27 cells in a tissue-treated T175 flask (Thermo Fisher Scientific) with 100 mL of supplemented DMEM. After 24 h, the media was removed, the cells were washed with sterile 1X PBS, and 50 mL of DMEM containing 1% P/S and no FBS was added. After an additional 48 h, the media was transferred to a centrifugal tube, centrifuged at 300 g for 5 min, and the supernatant was collected as the tumor conditioned media.

All cells were cultured at 37 °C in a humidified incubator with 5% CO_2_ and routinely tested for mycoplasma contamination through the University of Pennsylvania’s Cell Center Stockroom. Cells were counted using a Countess 3 automated cell counter (Thermo Fisher Scientific) using 1:1 cell solution and Trypan Blue Stain (Thermo Fisher Scientific) for cancer cells and no Trypan Blue Stain for monocytes and macrophages.

### In vitro assays

To evaluate luciferase expression, cells were plated at 20,000 cells per well in 100 µL of media in tissue culture treated white 96-well plates (Greiner) and were left to adhere for 16 h. Afterwards, the media was removed and replaced with the corresponding LNPs encapsulating FLuc mRNA at a concentration of 20 ng of mRNA per well in 100 µL of media. The cells incubated with the LNPs for 24 h. Luciferase expression was measured by removing the media, adding 50 µL of 1X reporter lysis buffer (Promega, Madison, WI) followed by 100 µL of luciferase assay substrate (Promega). The plate was then wrapped in aluminum foil and then shook on a plate shaker at 200 rpm for 10 min. Luminescence intensity was quantified using an Infinite 200 Pro plate reader (Tecan) and normalized based on the average intensity of the C12-200 LNP. Normalized luminescence is reported as mean ± SD of n = 3 biological replicates, each averaged from n = 4 technical replicates. Cellular toxicity of the LNPs were measured using the same cell setup as described above, where LNP dose depended on the specific experiment. After incubating the cells with LNPs for 24 h or 48 h, 100 µL of CellTiter-Glo (Promega) was added to each well, and then the plate was wrapped in aluminum foil and shook on a plate shaker at 200 rpm for 10 min. The luminescence corresponding to ATP concentration was quantified using an Infinite 200 Pro plate reader and normalized by dividing the luminescence from each group by the average luminescence signal of an untreated group. Cell viability percentage was reported as mean ± SD of n = 3 biological replicates, each averaged from n = 3 technical replicates.

Caspase-3/7 assays were performed using the AnaSpec (Fremont, CA, USA) SensoLyte Homogenous AMC Caspase-3/7 Fluorimetric Assay Kit. Tissue-treated black 96-well plates (Greiner) containing 20,000 cells per well were prepared as described above. After incubating LNPs at 5, 20, and 50 ng of mRNA per well for 24 h in a total volume of 150 µL, 50 µL of the Caspase-3/7 substrate solution was added to each well and the plate was wrapped in aluminum foil and shook on a plate shaker at 200 rpm for 1 min. Fluorescence corresponding to caspase-3/7 activity was quantified using an Infinite 200 Pro plate reader using an excitation wavelength of 354 nm and emission wavelength of 442 nm. Fluorescence was normalized by dividing the fluorescence signal from each group by the average fluorescence signal of an untreated group. RFU was reported as mean ± SD of n = 4 technical replicates.

Caspase-9 assays were performed using the Promega Caspase-Glo 9 Assay System. Tissue-treated white 96-well plates containing 20,000 cells per well were prepared as described above. After incubating LNPs at 50 ng of mRNA per well for 24 h, 100 µL of Caspase-Glo 9 Reagent was added to each well and the plate was wrapped in aluminum foil and shook on a plate shaker at 300 rpm for 2 min. The plate was then removed from the shaker and incubated at room temperature for 30 min. Luminescence corresponding to caspase-9 activity was quantified using an Infinite 200 Pro plate reader and normalized by dividing the luminescence from each group by the average luminescence signal of an untreated group. Relative luminescence was reported as mean ± SD of n = 3 technical replicates.

Mitochondrial permeability assays were performed using the APExBio (Houston, TX, USA) Mitochondrial Permeability Transition Pore Assay Kit. Tissue-treated black 96-well plates containing 20,000 CAL-27 cells per well were prepared as described above. LNPs were added to the cells at 5, 20, and 50 ng of mRNA per well for 24 h in a total volume of 100 µL. After 24 h, the Calcein AM working solution and fluorescence quenching working solution were added and the cells incubated at 37 °C for 30 min before being washed with dilution buffer. Fluorescence corresponding to mitochondrial permeability was quantified using an Infinite 200 Pro plate reader using an excitation wavelength of 485 nm and emission wavelength of 525 nm. Fluorescence was normalized by dividing the fluorescence signal from each group by the average fluorescence signal of an untreated group. RFU was reported as mean ± SD of n = 4 technical replicates.

Reactive oxygen species (ROS) assays were performed using the Thermo Fisher Scientific Total Reactive Oxygen Species Assay Kit (520 nm). Tissue-treated black 96-well plates containing 20,000 CAL-27 cells per well were prepared as described above. After adhering, the ROS Assay Stain was added to the cells and the cells incubated at 37 °C for 60 min. LNPs were added to the cells at 5, 20, and 50 ng of mRNA per well for 24 h in a total volume of 100 µL. After 24 h, fluorescence corresponding to ROS activity was quantified using an Infinite 200 Pro plate reader using an excitation wavelength of 480 nm and emission wavelength of 520 nm. Fluorescence was normalized by dividing the fluorescence signal from each group by the average fluorescence signal of an untreated group. RFU was reported as mean ± SD of n = 4 technical replicates.

Griess assays were performed using the Promega Griess Reagent System. TAMs were plated on non-tissue-treated black 96-well plates at density of 100,000 cells per well. Upon adhering for 16 h, LNPs were added at a dose of 100 ng of mRNA per well. After an additional 24 h, 50 µL of media from each well was transferred to a clear 96-well plate (Thermo Fisher Scientific). A nitrite standard curve was prepared according to the manufacturer’s instructions using concentrations ranging from 100 µM to 1.56 µM and included a blank well. Then, 50 µL of the Sulfanilamide solution was added to each well. The plate incubated for 5 min at room temperature away from light, before the addition of 50 µL of the NED solution to all wells. The plate incubated for another 5 min at room temperature protected from light. Absorbance corresponding to nitrite concentration was measured at 535 nm using an Infinite 200 Pro plate reader. Nitrite concentration was determined via a standard curve estimated from a LSLR. Nitrite concentration was reported as mean ± SD of n = 4 technical replicates.

### In vitro flow cytometry

For dose response curves, E10i-494 LNP was formulated with mCherry mRNA. Cancer cells were plated in tissue-treated 24-well plates (Greiner) while M0 macrophages and TAMs were plated in non-tissue-treated 24-well plates (Thermo Fisher Scientific) at cell densities of 250,000 cells per well. The cells adhered for 16 h, and then mCherry-containing E10i-494 was added to each well at mRNA doses ranging from 0.05 ng to 250 ng for cancer cells and 5 ng to 1,000 ng for M0 macrophages and TAMs. After 24 h of LNP incubation, media was removed, and the cells were washed with 1X PBS. The cancer cells were removed with 0.25% Trypsin-EDTA (Thermo Fisher Scientific) while the M0 macrophages and TAMs were removed with TrypLE Express (Gibco) followed by scraping with 40 cm cell scrapers (Greiner). After detachment, the oral cancer cells were diluted two-fold in supplemented DMEM while the M0 macrophages and TAMs were diluted two-fold in their corresponding media. The cells were centrifuged at 400 g for 5 min, where the supernatant was aspirated and the cells were resuspended in 1X PBS before being analyzed on a FACSymphony A5 SE cell analyzer (Becton Dickinson, Franklin Lakes, NJ). mCherry+ cells were reported as mean ± SD of n = 3 technical replicates.

For TAM phenotype experiments, TAMs were plated as described above and treated with LNPs at a dose of 250 ng of mRNA per well. After detaching and resuspension in 1X PBS, TAMs were extracellularly stained for 30 min with BV421-CD163 (BioLegend, San Diego, CA, USA), APC-CD38 (BioLegend), BV510-CD274 (BioLegend), and PE/Cy5-CD86 (BioLegend). The cells were then fixed, permeabilized, and stained intracellularly with AF488-Arg-1 (Thermo Fisher Scientific), PE-CD206 (Becton Dickson), APC/Cy7-CD68 (BioLegend), and AF594-p53 (Santa Cruz Biotechnology, Dallas, TX, USA) using the Cyto-Fast Fix/Perm Buffer Set (BioLegend) according to the manufacturer’s instructions. TAMs were resuspended in 1X PBS before being analyzed on a FACSymphony A5 SE cell analyzer. Median fluorescent intensity (MFI) was reported as mean ± SD of n = 5 technical replicates.

### Murine tumor burden studies with CAL-27 cells

Nu/J male mice of 6-8 weeks old with an average weight of 25 g were purchased from Jackson Laboratory (Bar Harbor, ME). Animals were housed in a barrier facility with air humidity 40%–70%, ambient temperature of 22 ± 2 °C, and 12 h dark/light cycle. The mice were inoculated on the right flank with two million CAL-27 cells suspended in 1X PBS and Matrigel (Corning, Corning, NY, USA). After two weeks, for biodistribution studies LNPs encapsulating FLuc mRNA were intratumorally injected into the tumors at a dose of 0.1 mg of mRNA per kg of body weight (mpk) using a total volume of 20 µL. After 6 h, the mice were intraperitoneally injected with D-luciferin (0.2 mL, 15 mg/mL; Biotium, Fremont, CA, USA). After 5 min, the mice were euthanized by CO_2_ asphyxiation followed by cervical dislocation. Then, the heart, lungs, liver, kidneys, spleen, and tumor were dissected and imaged using an In Vivo Imaging System (IVIS; Revvity, Waltham, MA, USA). Total flux was quantified by the Living Image Software 4.7.3 (PerkinElmer) by placing rectangular region of interests (ROI) around the organ images, keeping the same ROI sizes among each organ. Total flux was reported as mean ± SD of n = 3 biological replicates.

For tumor burden, mice were inoculated as described above, where each mouse received injections on the left and right flanks. Two weeks after inoculation, LNPs were injected intratumorally four times over the course of ten days at 0.1 mpk using a total volume of 20 µL in each tumor. Tumor volume was monitored using calipers and calculated by 0.5 × width^2^ × height. After eighteen days from the first dose, the mice were sacrificed, and tumors were extracted for further analysis by immunohistochemistry (IHC) for human p53 and human PUMA expression. Slides were prepared by the Histotechnology Facility at the Wistar Institute. Images were acquired using an EVOS M7000 Imaging System (Thermo Fisher Scientific). Fiji/ImageJ v1.54j was utilized to process IHC slides through the plug-in Colour Deconvolution2, selecting “H DAB” to separate out stained protein of interest. Percent area is measured from *n* = 10 images with a solid representing the median and dashed lines representing quartiles

### Murine tumor burden studies with MOC-2 cells

C57BL/6J female mice of 6–8 weeks old with an average weight of 20 g were purchased from Jackson Laboratory. The fur on the right flank was shaved and then the flank was inoculated with 2×10^5^ MOC-2 cells suspended in 0.1 mL of 1X PBS containing 25% Matrigel. After seven days, LNPs were injected intratumorally four times over the course of ten days at 0.5 mpk using a total volume of 20 µL. After a total of sixteen days from the first LNP injection, the mice were sacrificed, and the tumors were isolated for IHC and flow cytometry. For IHC, tumor tissues were fixed with 4% PFA (MilliporeSigma), and paraffin-embedded 5 μm sections were deparaffinized with xylene (MilliporeSigma), rehydrated with 200 proof ethanol (Thermo Fisher Scientific), and heated in 10 mmol/L sodium citrate buffer (MilliporeSigma) at pH 6.0 for antigen retrieval. After blocking with 2.5% goat serum (Thermo Fisher Scientific) in 1X PBS, the sections were incubated for 16 h at 4 °C with primary antibodies, then detected using a universal immunoperoxidase ABC kit. All the sections were counterstained with hematoxylin. Images were acquired and quantified as described above. Percent area is measured from *n* = 8 images with a solid representing the median and dashed lines representing quartiles For flow cytometry, the tumors were diced and incubated in digestion medium containing DNase I (New England Biolabs; Ipswich, MA), collagenase IV (Thermo Fisher Scientific), and dispase II (Fisher Scientific) for 1 hour at 37 ºC. Afterwards, the solution was passed through a 70 µm filter (Corning) into 15 mL centrifugal tubes (Corning). The tubes were centrifuged at 500 g for 5 min. The supernatant was removed and the cells were resuspended in ACK Lysis Buffer (Thermo Fisher Scientific) on ice for 5 min. Afterwards, the solution was diluted 2.5-fold in FACS buffer containing 0.6% BSA, 25 mM 4-(2-hydroxyethyl)-1-piperazineethanesulfonic acid (HEPES; Thermo Fisher Scientific), and 1 mM ethylenediaminetetraacetic acid (EDTA; Thermo Fisher Scientific) in 1X PBS. The cells were then centrifuged at 500 g for 5 min, and the supernatant was aspirated. Then, the cells were stained with Live/Dead Zombie Aqua (BioLegend) according to the manufacturer’s instructions, followed by extracellular staining with BV605-CD163 (BioLegend), eFluor450-CD86 (Thermo Fisher Scientific), BV650-F4/80 (BioLegend), APC/Fire750-CD11b (BioLegend), and BUV395-CD45 (Becton Dickinson). The cells were then fixed, permeabilized, and stained intracellularly with PE/Fire700-CD206 (BioLegend) and AF488-Arg-1 (Thermo Fisher Scientific) using the Cyto-Fast Fix/Perm Buffer Set (BioLegend) according to the manufacturer’s instructions. The cells were resuspended in FACS buffer before being analyzed on a FACSymphony A5 SE cell analyzer. Median fluorescent intensity (MFI) was reported as mean ± SD of n = 4 biological replicates.

### Statistical & reproducibility

All statistical analysis was performed in GraphPad Prism Version 10.5.0 (GraphPad Software, Inc, La Jolla, USA). All tests of significance were performed at a significance level of α = 0.05. For experiments with one variable where multiple technical or biological replicates were performed, one-way analyses of variance (ANOVAs) with post hoc Holm-Šídák correction for multiple comparisons were used to compare responses across treatment groups. For experiments that measured two variables with more than two treatment groups, a two-way ANOVA with post hoc Holm-Šídák correction for multiple comparisons were used to compare responses across treatment groups. For experiments that measured two variables with two treatment groups in each variable, two-tailed multiple unpaired t tests with Holm-Šídák correction for multiple comparisons were used to compare responses across treatment groups. For mouse survival, a Kaplan-Meier estimator was employed with post hoc Holm-Šídák correction for multiple comparisons to compare survival curves across treatment groups. All data are presented as mean ± standard deviation unless otherwise reported. No statistical method was used to predetermine sample size. No data were excluded from the analyses. The experiments were not randomized. The investigators were not blinded to allocation during experiments and outcome assessment.

## Data availability

All relevant data supporting the findings of this study are available within the paper and Supplementary Information.

## Code availability

No original code was developed for this manuscript.

## Acknowledgements

M.S.P. acknowledges support from the Center for Innovation & Precision Dentistry (CiPD) at the University of Pennsylvania. The authors thank Emily Cento, Zhilin Chen, Max A. Eldabbas, and Emileigh Maddox of the Human Immunology Core and the Division of Transfusion Medicine and Therapeutic Pathology at the Perelman School of Medicine at the University of Pennsylvania for providing de-identified T cells that were purified from healthy donor apheresis using StemCell RosetteSep™ kits. Figure schematics were created with BioRender.

## Author Contributions

M.S.P., Q.Z., A.D.L., and M.J.M. designed the experiments. M.S.P., J.J.L., Q.Z., K.P., S.S., H.M.Y., R.A.J., B.N.H., and S.V.T. performed the experiments. M.S.P., J.J.L., Q.Z., K.P., and S.V.T. analyzed the data. M.S.P. wrote the manuscript. Q.Z., A.D.L., M-G.A., D.W. edited the manuscript. All authors discussed the results and commented on the manuscript.

