## Supplementary Information for "Lipid nanoparticle co-delivery of mRNA and a small molecule drug for oral cancer chemoimmunotherapy"

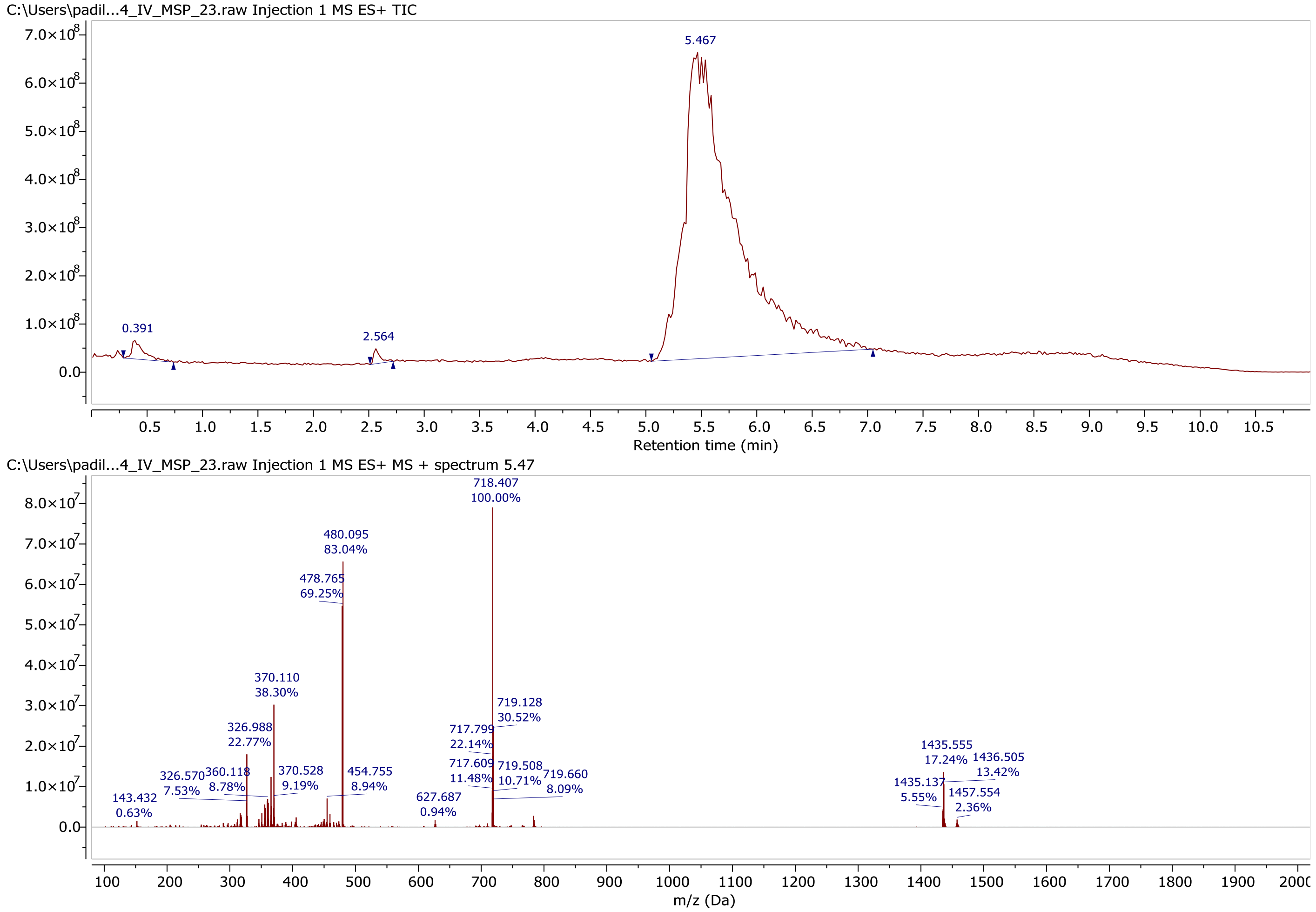


**Supplementary Figure 1. LC-MS trace of E10i-494.** Electrospray ionization mass spectrometry calculated for E10i-494, 1435.47; found 1435.56. Purity: 97.94%

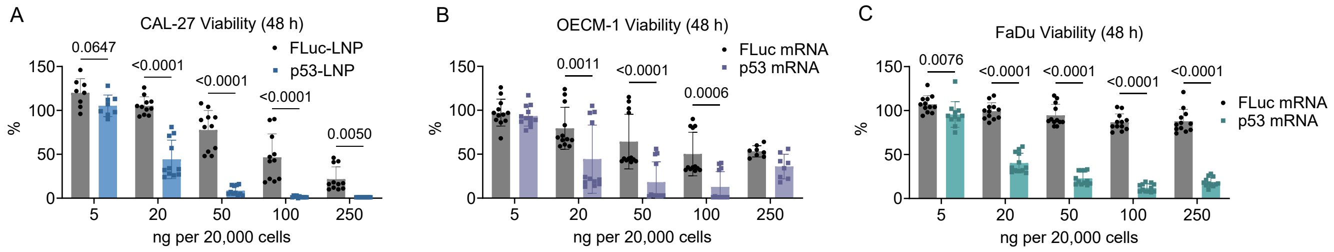


**Supplementary Figure 2. In vitro toxicity of p53-mRNA-loaded LNPs after 48 h. A–C,** E10i-494 was formulated with FLuc mRNA (FLuc-LNP) and human p53 mRNA (p53-LNP). The two LNPs were incubated in **(A)** CAL-27, **(B)**, OECM-1, and **(C)** FaDu cells at increasing dosages per 20,000 cells, and after 48 h, viability was measured. Viability is reported as mean ± SD of $n$ = 9–12. Multiple unpaired t test with *post hoc* Holm–Šídák correction for multiple comparisons was used to compare FLuc-LNP to p53-LNP.

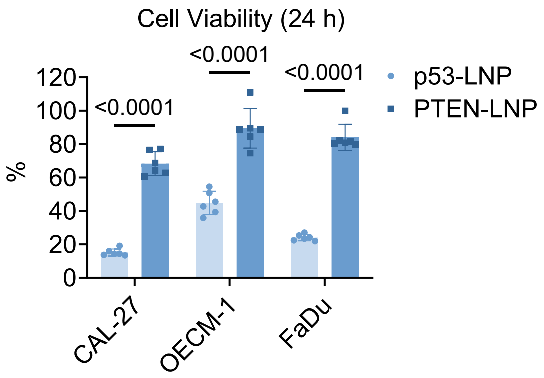


**Supplementary Figure 3. Comparison between LNPs encapsulating PTEN mRNA and p53 mRNA.** E10i-494 encapsulating human p53 mRNA (p53-LNP) and PTEN mRNA (PTEN-LNP) were dosed in CAL-27, OECM-1, and FaDu at 20 ng per 20,000 cells, and after 24 h, cellular viability was measured. Viability is reported as mean ± SD of $n$ = 6. Multiple unpaired t test with *post hoc* Holm–Šídák correction for multiple comparisons was used to compare p53 mRNA to PTEN mRNA.

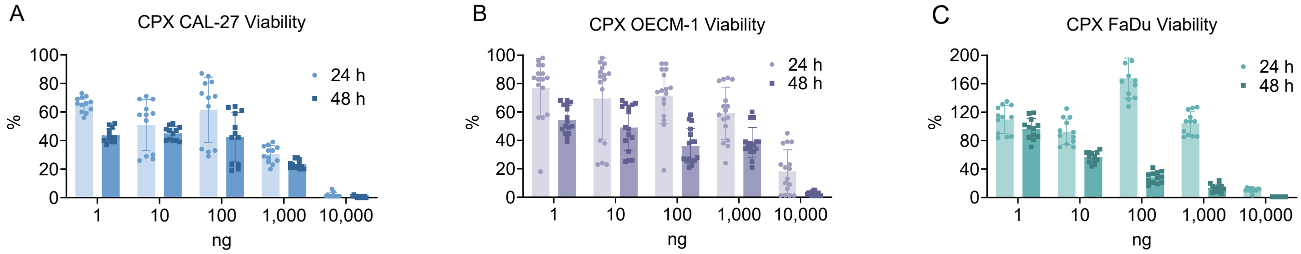


**Supplementary Figure 4. CPX chemotherapeutic efficacy in OSCC lines.** Free CPX was incubated with **(A)** CAL-27, **(B)**, OECM-1, and **(C)** FaDu cells at increasing doses per 20,000 cells. After 24 and 48 h, cellular viability was measured. Viability is reported as mean ± SD of $n$ = 12.

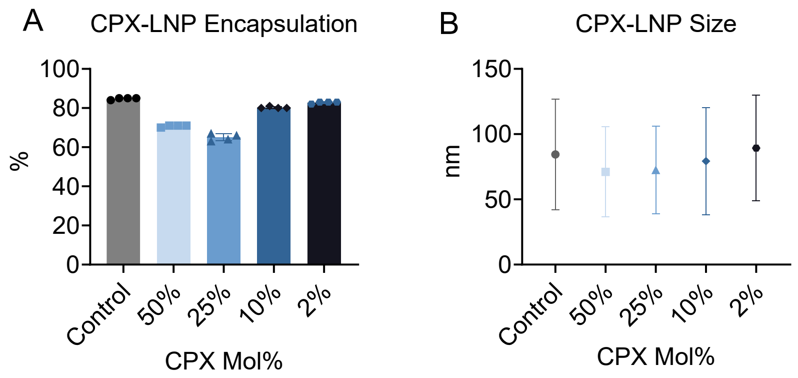


**Supplementary Figure 5. Physicochemical characteristics of CPX-loaded LNPs. A,** Relative encapsulation efficiency determined by a RiboGreen assay of E10i-494 LNPs encapsulating FLuc-mRNA and increasing mol% of CPX, where control refers to the LNP with no CPX. Percent encapsulation is reported as mean ± SD of $n$ = 4. **B,** Hydrodynamic radii determined by dynamic light scattering of E10i-494 LNPs encapsulating FLuc-mRNA and increasing mol% of CPX. Hydrodynamic radii are reported as mean ± SD of $n$ = 3.

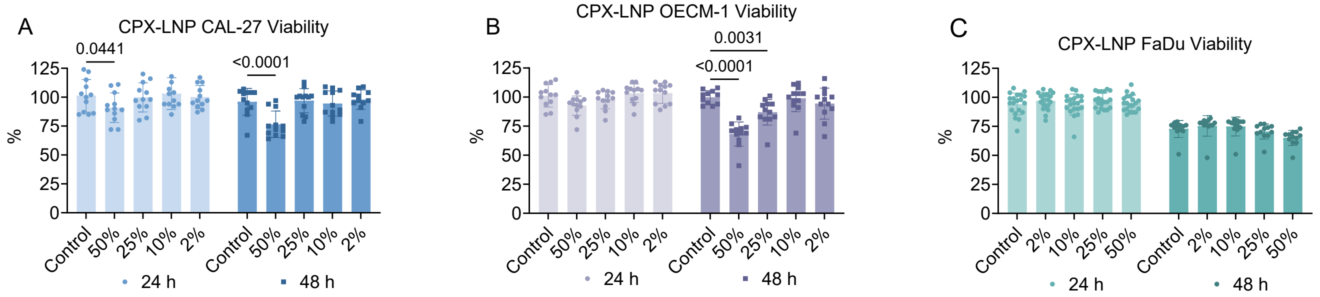


**Supplementary Figure 6. CPX-LNP toxicity in OSCC lines.** E10i-494 encapsulating increasing mol% of CPX were dosed in **(A)** CAL-27, **(B)** OECM-1, and **(C)** FaDu at 20 ng per 20,000 cells, and after 24 and 48 h, cellular viability was measured. Control formulation has no CPX. Viability is reported as mean ± SD of $n$ = 12. Two-way ANOVA with *post hoc* Holm–Šídák correction for multiple comparisons was used to compare the control LNP against all other formulations for each time point.

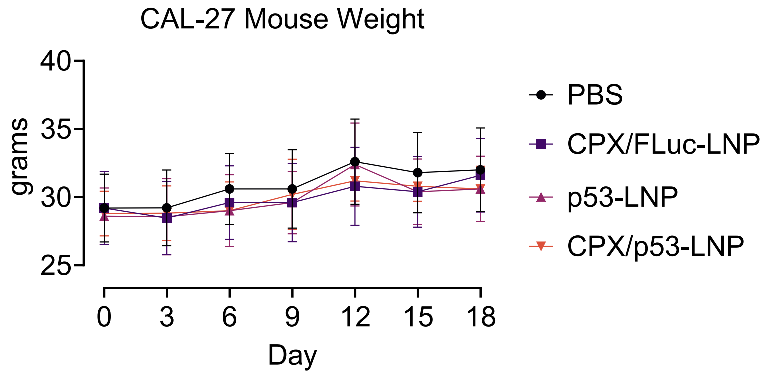


**Supplementary Figure 7. Mouse weights for CAL-27 xenograft study.** Nu/J mice were inoculated with two million CAL-27 cells on each flank, and after two weeks, LNPs were administered intratumorally at an mRNA dose of 0.1 mpk four times over ten days on each tumor. Mouse weight was monitored from the initial dose on day 0 until day 18. Weight is reported as mean ± SD of $n$ = 5.

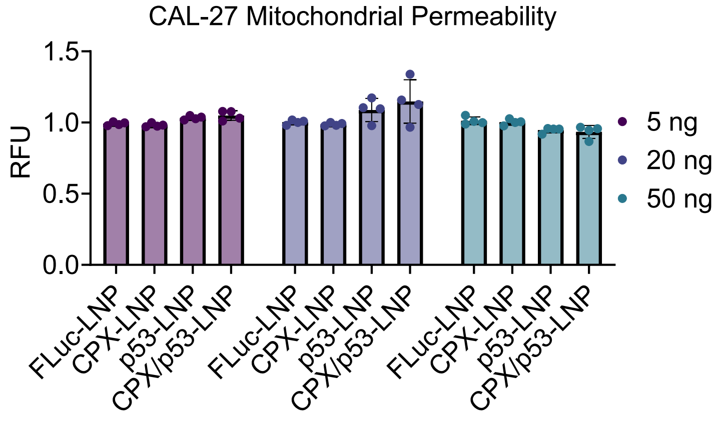


**Supplementary Figure 8. Mitochondrial permeability assay.** E10i-494 LNPs containing FLuc mRNA (FLuc-LNP), 50 mol% CPX (CPX-LNP), human p53 mRNA (p53-LNP), and 50 mol% CPX and human p53 (CPX/p53-LNP) were dosed in CAL-27 cells at 5, 20, and 50 ng per 20,000 cells. After 24 h, a fluorescence-based mitochondrial permeability assay was performed, where fluorescence was calculated relative to untreated cells. Relative fluorescent units (RFU) are reported as mean ± SD of $n$ = 4. Two-way ANOVA with *post hoc* Holm–Šídák correction for multiple comparisons was used to compare individual LNPs for each dose.

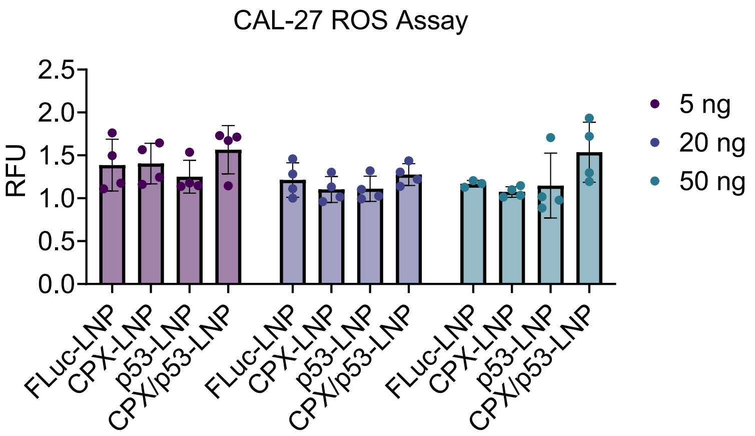


**Supplementary Figure 9. ROS assay.** E10i-494 LNPs containing FLuc mRNA (FLuc-LNP), 50 mol% CPX (CPX-LNP), human p53 mRNA (p53-LNP), and 50 mol% CPX and human p53 (CPX/p53-LNP) were dosed in CAL-27 cells at 5, 20, and 50 ng per 20,000 cells. After 24 h, a fluorescence-based reactive oxygen species (ROS) assay was performed, where fluorescence was calculated relative to untreated cells. Relative fluorescent units (RFU) are reported as mean ± SD of *n* = 4. Two-way ANOVA with *post hoc* Holm–Šídák correction for multiple comparisons was used to the control against all other formulations.

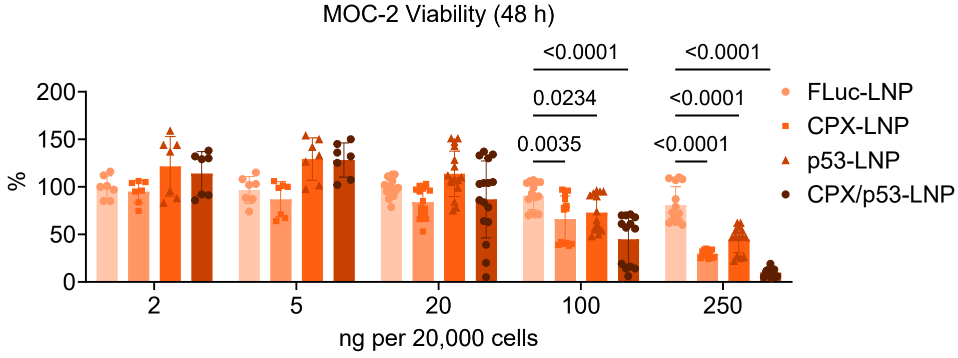


**Supplementary Figure 10. MOC-2 viability after 48 h of therapeutic mRNA treatment.** E10i-494 LNPs containing FLuc mRNA (FLuc-LNP), 50 mol% CPX (CPX-LNP), mouse p53 mRNA (p53-LNP), and 50 mol% CPX and mouse p53 (CPX/p53-LNP) were incubated in MOC-2 cells at increases doses per 20,000 cells, and after 48 h, viability was measured. Cellular viability is reported as mean ± SD of *n* = 8 for 2 ng and 5 ng doses and *n* = 12 for all other doses. Two-way ANOVA with *post hoc* Holm–Šídák correction for multiple comparisons was used to the FLuc-LNP against all other formulations for each dose.

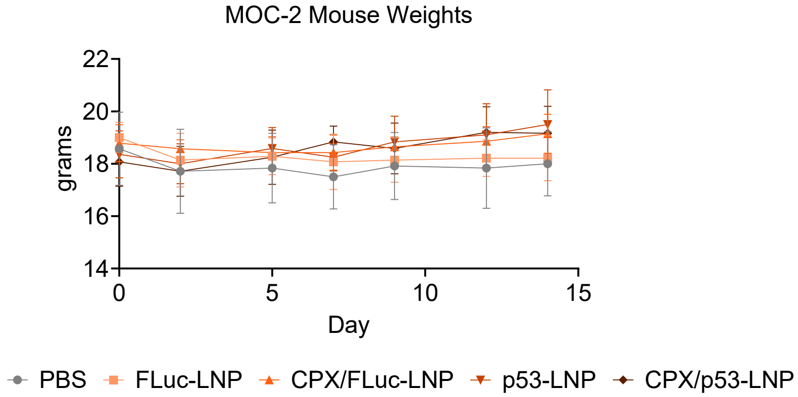


**Supplementary Figure 11. Mouse weights for MOC-2 survival study.** C57BL/6J mice were inoculated with 20,000 MOC-2 cells on the right flank, and after seven days, LNPs were administered intratumorally at an mRNA dose of 1.0 mpk four times over ten days. Mouse weight was monitored from the initial dose on day 0 until day 14. Weight is reported as mean ± SD of *n* = 6 for PBS and *n* = 5 for all other groups
